# Generation and characterisation of mouse models of Duchenne Muscular Dystrophy (DMD)

**DOI:** 10.1101/2024.08.11.607106

**Authors:** Jayshen Arudkumar, Yu Chinn Joshua Chey, Sandra G Piltz, Paul Q Thomas, Fatwa Adikusuma

## Abstract

CRISPR-Cas9 gene-editing technology has revolutionised the creation of precise and permanent modifications to DNA, enabling the generation of diverse animal models for investigating potential treatments. Here, we provide a protocol for the use of CRISPR-Cas9 to create murine models of Duchenne Muscular Dystrophy (DMD) along with a step-by-step guide for their phenotypic and molecular characterisation. The experimental procedures include CRISPR microinjection of embryos, molecular testing at the DNA, RNA, and protein levels, forelimb grip strength testing, immunostaining and serum creatine kinase (CK) testing. We further provide suggestions for analysis and interpretation of the generated data, as well as the limitations of our approach. These protocols are designed for researchers who intend to generate and use mouse models to study DMD as well as those seeking a detailed framework of phenotyping to contribute to the broader landscape of genetic disorder investigations.

## Background

Duchenne Muscular Dystrophy (DMD) is a progressive and fatal muscle-wasting disorder, primarily affecting boys, with an estimated incidence of 1 in 5,000 births. There are a wide range of causative loss-of-function mutations in the X-linked *DMD* gene that encodes for the dystrophin protein (1, 2). The most prevalent are large deletions, of which 80% occur within mutational hotspots within exons 2-20 and 45-55 (4). The absence of muscle dystrophin protein results in muscle wasting and damage, leading to the loss of independent ambulation during the pre-teen years. This condition becomes more severe with age, resulting in premature mortality due to progressive wasting of respiratory muscles and development of dilated cardiomyopathy (3). Several therapies have been developed to combat this disorder, although they are transient in effect with limited functional benefit on patients. As such, there remains no curative treatment option present for DMD (3).

The CRISPR-Cas9 gene editing technology utilises single guide RNA (sgRNA) coupled with Cas9 endonuclease to generate precise double-stranded breaks (DSBs) at targeted DNA sequences (5). These breaks prompt the cell’s innate DNA damage repair mechanisms, predominantly the error-prone non-homologous end joining (NHEJ) pathway active throughout the cell cycle, often resulting in insertions or deletions (InDels) surrounding the break site (9). In instances where two gRNAs are positioned closely on the same chromosome, targeted removal of the intervening region can occur. CRISPR-Cas9 technology signifies a groundbreaking advancement in generating genetically modified animal models (6). This innovative approach provides a swift and cost-effective method to engineer genetic modifications for animal model creation. Over the past decade, CRISPR-Cas9 has gained global traction in laboratories, transforming the landscape of genetically modified animal model generation with unparalleled speed and precision (6, 10).

The use of CRISPR-Cas9 technology has been instrumental in creating bespoke mouse models for Duchenne Muscular Dystrophy (DMD), serving as indispensable tools to explore both the molecular intricacies and physical traits of this condition (10). These models offer significant insights into disease mechanisms and provide physiologically-relevant platforms for therapeutic development. Moreover, CRISPR-Cas9 has emerged as a potential therapeutic approach for DMD, with numerous studies focusing on developing CRISPR therapies for this condition. Therefore, establishing robust, reliable, and easily accessible protocols for generating DMD mouse models and assessing DMD phenotypes would be beneficial. This could address the current issue of fragmented protocols scattered across multiple publications, thereby simplifying the workflow for researchers. In this comprehensive protocol, we delineate the step-by-step process for generating DMD mouse models. Specifically, our focus was on creating exon Δ51 and exon Δ52 mouse models, which are prevalent frameshift-inducing mutations in DMD patients. Our protocol also offers a streamlined, detailed procedure for the thorough physical and molecular characterisation of dystrophic pathology within these models. By following these precise protocols, researchers can conduct meticulous examinations, establishing a robust foundation for further investigations into disease mechanisms and potential therapeutic interventions.

## Materials and Procedure

The protocols described in this article are published on protocols.io (see links below) and are included for printing as supporting information file 1 with this article, updated on 29 January 2024.

There are 9 separate protocols with each of the reserved DOIs listed here:

In vitro Transcription protocol: https://www.protocols.io/view/in-vitro-transcription-ivt-of-guide-rnas-for-cytop-c7jnzkme

Lateral Tail Vein Bleeding protocol: https://www.protocols.io/view/lateral-tail-vein-bleeding-for-blood-collection-in-c7yrzpv6

Post-mortem Tissue protocol: https://www.protocols.io/view/postmortem-mouse-tissue-processing-necropsy-c7ywzpxe

DNA Extraction protocol: https://www.protocols.io/view/genomic-dna-extraction-and-genotyping-pcr-on-mouse-c8d9zs96

Immunofluorescence Staining protocol DOI: https://www.protocols.io/view/scientific-protocol-for-immunofluorescence-analysi-c7yyzpxw

H&E Staining protocol: https://www.protocols.io/view/scientific-protocol-for-h-amp-e-staining-in-heart-c7yzzpx6

RNA Extraction protocol: https://www.protocols.io/view/rna-extraction-and-rt-pcr-on-mouse-tissues-c732zqqe

Western Blot protocol: https://www.protocols.io/view/scientific-protocol-for-western-blotting-c736zqre

Grip strength testing protocol: https://www.protocols.io/view/forelimb-grip-strength-testing-c737zqrn

### Generation of DMD mouse models

All experiments involving animal use were approved by the South Australian Health & Medical Research Institute (SAHMRI) Animal Ethics Committee. To produce the guide RNAs, we first employed in-silico screening for the selection of intron-targeting guides using CRISPOR (12). Oligonucleotide pairs containing the gRNA spacer sequences were cloned into the pSpCas9(BB)- 2A-Puro (PX459) V2.0 (Addgene #62988) vector (7) using *Bbs*I-mediated Golden-Gate cloning method. These vectors were then PCR amplified to produce in vitro transcription (IVT) templates. PCR was performed using the primers listed in S1 Table. In brief, the forward primer contained both T7 promoter and gRNA sequences, while the reverse primer contained the tracrRNA sequence. The PCR amplicon was purified using the QIAquick PCR Purification Kit (Qiagen). The purified PCR products were in-vitro transcribed using the HiScribe T7 Quick High Yield RNA Synthesis Kit (NEB). SpCas9 nuclease mRNA was produced by IVT of Xho-linearised CMV/T7-hCas9 (Toolgen). SpCas9 nuclease mRNA and gRNA were purified using RNeasy Mini Kit (Qiagen). To generate the mouse models, SpCas9 nuclease mRNA (100 ng/μl) and two gRNAs (each at 50 ng/μl) were injected into the cytoplasm of C57BL/6J zygotes using a FemtoJet microinjector (11). Surviving zygotes were transferred into pseudo-pregnant females. Founder males and females with the desired hemizygous knockout and homozygous knockout genotype were then set up for breeding.

### Genomic DNA extraction and PCR

Mouse genomic DNA was extracted from the ear notch tissues using the High Pure PCR Template Preparation Kit (Roche), according to the manufacturer’s protocol. PCR reactions were set up with Phusion DNA Polymerase (New England Biolabs). PCR primers used in this study can be found in S2 Table.

### RNA, cDNA generation, RT PCR and qPCR

Total RNA was extracted from mouse tissues using Trizol reagent (Invitrogen) followed by the RNEasy mini extraction kit (Qiagen). RNA was quantitated using the Nanodrop One Microvolume Spectrophotometer (ThermoFisher Scientific). For cDNA generation, 1-2 μg of total RNA was used as template for first-strand cDNA synthesis using the High-Capacity cDNA Reverse Transcription Kit (Applied Biosystems) as per the manufacturer’s instructions. For SYBR Green RT-qPCR, expression levels of genes were determined using primers spanning the exon-exon boundaries. All 15 μl RT-qPCR reactions contained 1 μl cDNA, 1x Fast SYBR Green Real-Time PCR master mix (Applied Biosystems) and 0.5 μM forward and reverse primer mix. Actin Beta (*Actb*) was used as a housekeeping gene to normalise data. Data was collected and analysed using the QuantStudio Real-Time PCR software (Applied Biosystems).

### Dystrophin Western blot analysis

For Western blot of skeletal or heart muscle, tissues were homogenised in disruption buffer (15% SDS, 75 mM Tris HCl pH 6.8) and protease inhibitor cocktail (Roche), using green magna lyser tubes (Roche) at 6500RPM for 2 cycles of 20 s. Samples were then centrifuged at 4°C at max speed for 10 minutes before isolating the supernatant. Protein concentration was determined by a BCA Protein Assay Kit (ThermoFisher Scientific), and 50 μg of total protein was loaded onto a NuPAGE™ 3 to 8% Tris-Acetate protein gel (ThermoFisher Scientific). Gels were run at 100V for 15 minutes, followed by 120V for 1 hour and 45 minutes. This was followed by a 10-min transfer using the ‘HIGH MW’ program in the Trans-Blot Turbo Transfer System (Bio-Rad). The blot was blocked using a 10% Skim milk-PBST for 1 hour and then incubated with mouse anti-dystrophin antibody (MANDYS8, Sigma-Aldrich, D8168) at 4°C overnight (13, 14). The blot was washed and incubated with goat anti-mouse antibody conjugated with horseradish peroxidase (HRP) (Bio-Rad Laboratories) at room temperature for 1 hour. The blot was developed using Supersignal West Femto kit (ThermoFisher Scientific). The loading control was determined by blotting with mouse anti-vinculin antibody (Sigma-Aldrich, V9131).

### Histological analysis of muscles

Skeletal muscles from wild-type (WT) and DMD mice were individually dissected and cryo-embedded in 10% Gum Tragacanth (Sigma-Aldrich) on small cork blocks. All embeds were snap frozen in isopentane supercooled in liquid nitrogen. The resulting blocks were stored at −80°C prior to sectioning. 10 μm transverse sections of the skeletal muscle and frontal sections of the heart were prepared and air-dried prior to staining. H&E staining was performed with some modifications to the established TREAT-NMD standard operating procedure DMD_M.1.2.007 and a separate study by the Yokota lab (24). Dystrophin immunohistochemistry was performed using MANDYS8 monoclonal antibody (Sigma-Aldrich) with modifications to the manufacturer’s instructions. In brief, cryostat sections were thawed and washed with 4% PFA before being immersed in permeabilisation solution for 10-min. Next, slides were treated with blocking solution and incubated with 1x Mouse-on-Mouse IgG Blocking Solution (ThermoFisher Scientific). Slides were then washed and treated with MANDYS8 diluted 1:100 in blocking solution. Following overnight primary antibody incubation at 4°C, sections were washed, incubated with Donkey anti-Mouse IgG (H+L) Alexa Fluor™ 594, washed, and nuclei were counterstained with ProLong™ Gold Antifade Mountant with DAPI (ThermoFisher Scientific) prior to coverslipping.

### Tail vein bleeding, Serum CK and Grip strength Analysis

Blood was isolated from the tail vein of male mice at P32. A skin numbing cream (EMLA) was applied to the tail of mice, followed by warming the animal in a mini thermacage (Datesand) at 37°C for 10 minutes. An incision was made over the lateral vein area of mice under restraint, and blood was collected into a microvette tube. The collected blood samples were allowed to clot at room temperature for 30 minutes before being spun down at 2000 RCF for 10 minutes at 4°C. The resulting colourless serum was frozen at -80°C for subsequent analysis. Serum creatine kinase levels were measured by a third-party laboratory (Gribbles Veterinary Pathology, Australia), using the ADVIA® Chemistry Creatine Kinase (CK_L) reagent with the ADVIA Chemistry 1800 System. The output values were presented as U/L. For forelimb grip strength testing, each mouse was weighed prior to the test. A force transducer (ANDILOG) equipped with a grasping grid (SDR Scientific) was set up to measure the peak force from each mouse applied during a pull. Pulling the mouse away from the grid once their front paws have clasped onto it revealed the highest force applied. Each mouse performed three series of pulls across four sets for a total of 12 pulls. Each set of three pulls was followed by a resting period of at least one minute. The maximum grip strength was determined by averaging the three highest values and normalising for body weight. The protocol listed here was adapted from the TREAT-NMD standard operating procedure SOP DMD_M.2.2.001 for grip strength testing.

### Statistics

All data are presented as mean ± SEM. One-way ANOVA with Tukey’s multiple comparisons were performed for comparison between the respective two groups (WT and ΔEx51 DMD mice, WT and ΔEx52 DMD mice). Data analyses were performed with statistical software (GraphPad Prism 10 Software, San Diego, CA, USA). *P* values less than 0.05 were considered statistically significant. Sample sizes for each experiment are stated in the figure legends and represent independent biological replicates in most instances.

### General notes and troubleshooting

Conducting the procedures and analyses outlined in this protocol entails several challenges and potential pitfalls (Table 1). One challenge is the difficulty in obtaining mouse founders with the desired deletion. This may arise due to ineffective Cas9-gRNA cutting, often caused by low-quality Cas9 mRNA or gRNAs used for zygote microinjection. To address these issues, we recommend producing a new batch of Cas9 mRNA or gRNAs. Alternatively, Cas9 mRNA can be obtained from a reputable supplier, or commercial Cas9 protein can be utilised. Additionally, reviewing guide RNA designs and sequences of purchased oligos can help ensure accurate design and oligo quality. Consideration should also be given to changing gRNA target sequences if certain gRNAs are not cutting efficiently, possibly due to the presence of gRNA-blocking motifs. The failure to obtain the desired founder can also be attributed to the microinjection operator. It is desirable that the microinjection is performed by a skilled and experienced microinjectionist.

**Table 1:**
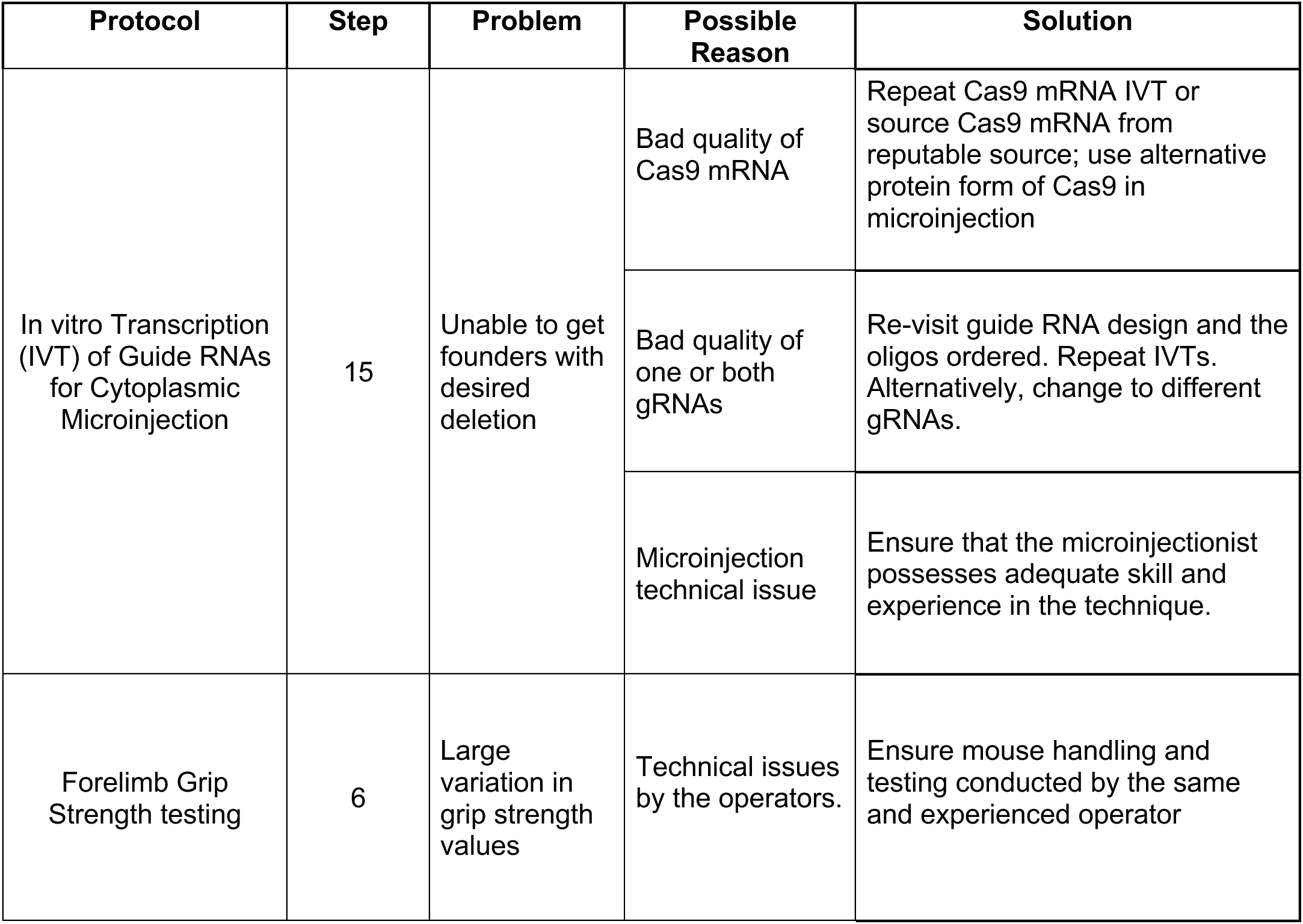
Protocol troubleshooting guide: Common problems, causes and solution.

The in vivo grip strength testing presents a certain level of variability and requires considerable time investment to achieve consistent reliable results. The variability may stem from several factors that include fatigue or habituation of mice to the testing procedure, behavioural changes in mice with respect to the testing environment or handling and inconsistencies in data recording by the operator. To obtain reliable and reproducible results, multiple assessments were done by the same operator, where we assayed a maximum of 3 determinations per mouse with 1-min rest intervals between each pull. The TREAT-NMD SOP DMD_M.2.2.001 elucidates the various biases that could arise irrespective of operator proficiency (29). Fatigue, for instance, stands out as a critical factor that may introduce confounding elements into sample readings. In addressing this concern, Aartsma-Rus and colleagues recommend the inclusion of fatigue calculations across each set of readings, providing a methodological approach to mitigate potential sources of variability and ensuring a more robust evaluation of grip strength dynamics (28, 29).

## Expected results

### Generation of DMD exon deletion mouse models

We employed CRISPR-Cas9 to generate two distinct mouse models, harbouring targeted deletions of either exon 51 (ΔEx51) or exon 52 (ΔEx52) of the *Dmd* gene (Fig 1). Firstly, we selected two gRNAs for each model that targeted the introns flanking the exon of interest, with cut site distances of 679 bp for the exon 51 deletion and 364 bp for the exon 52 deletion model to facilitate efficient founder screening via PCR (Table 2, Fig 2A). Standard PCR genotyping with primers flanking the cut sites identified a subset of G0 (founder) pups displaying smaller amplicons, indicating the expected fragment deletions resulting from the dual CRISPR targeting. An average of 67% of G0 pups carried at least one copy of the desired deletions. Subsequent Sanger sequencing of these PCR products confirmed the correct removal of the target exons (Figure 2B). These founders were mated with WT mice to establish colonies.

**Figure 1:**
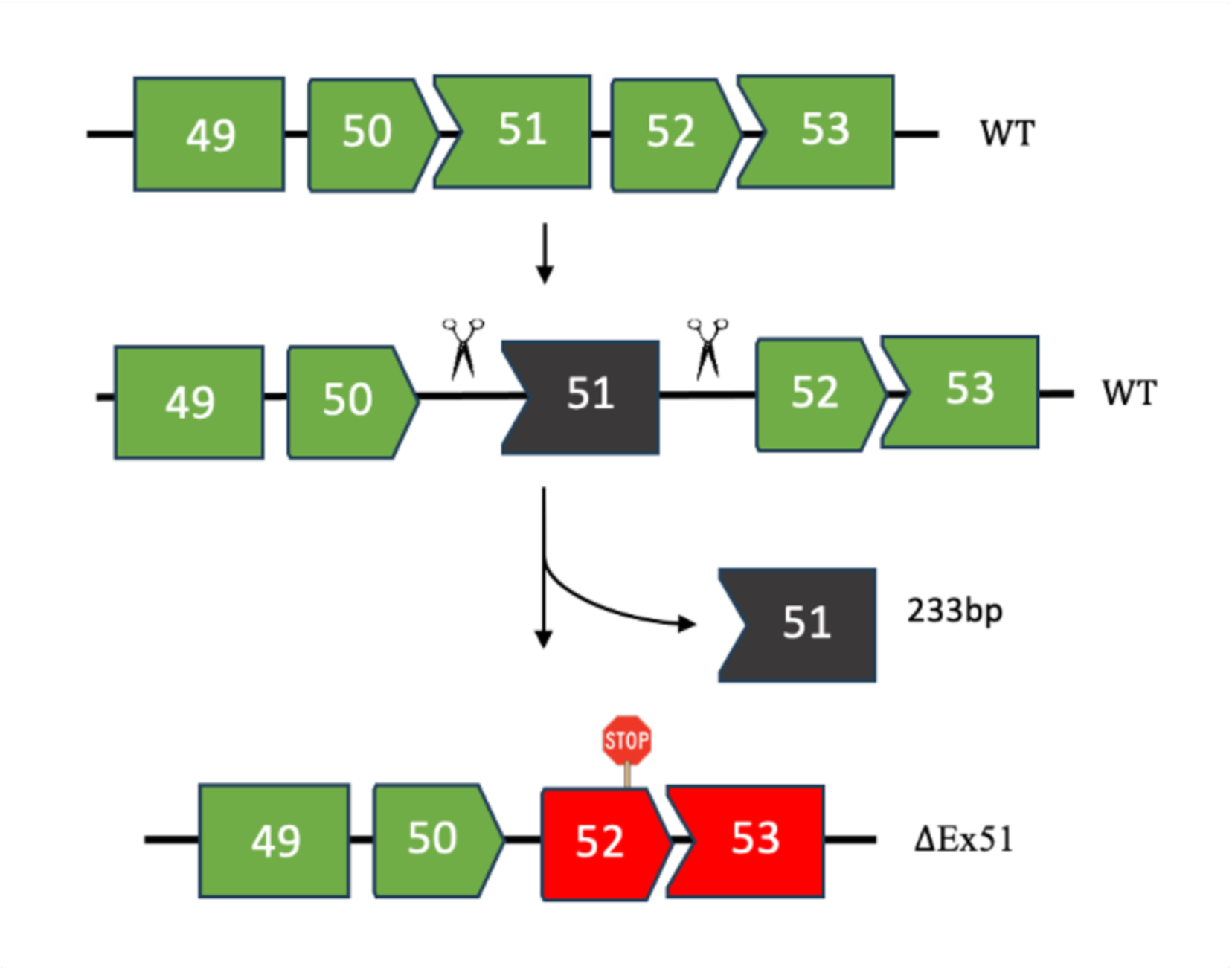
CRISPR-Cas9 editing strategy for generation of mice with exon 51 deletion (ΔEx51). Exon 52 (red) is out of frame with exon 50. The same intron-targeting strategy was applied in the context of generating an exon 52 deletion (ΔEx52) mouse model. In the latter case, intron 51 and intron 52 were cut to result in Exon 53 being out of frame with Exon 51.

**Figure 2:**
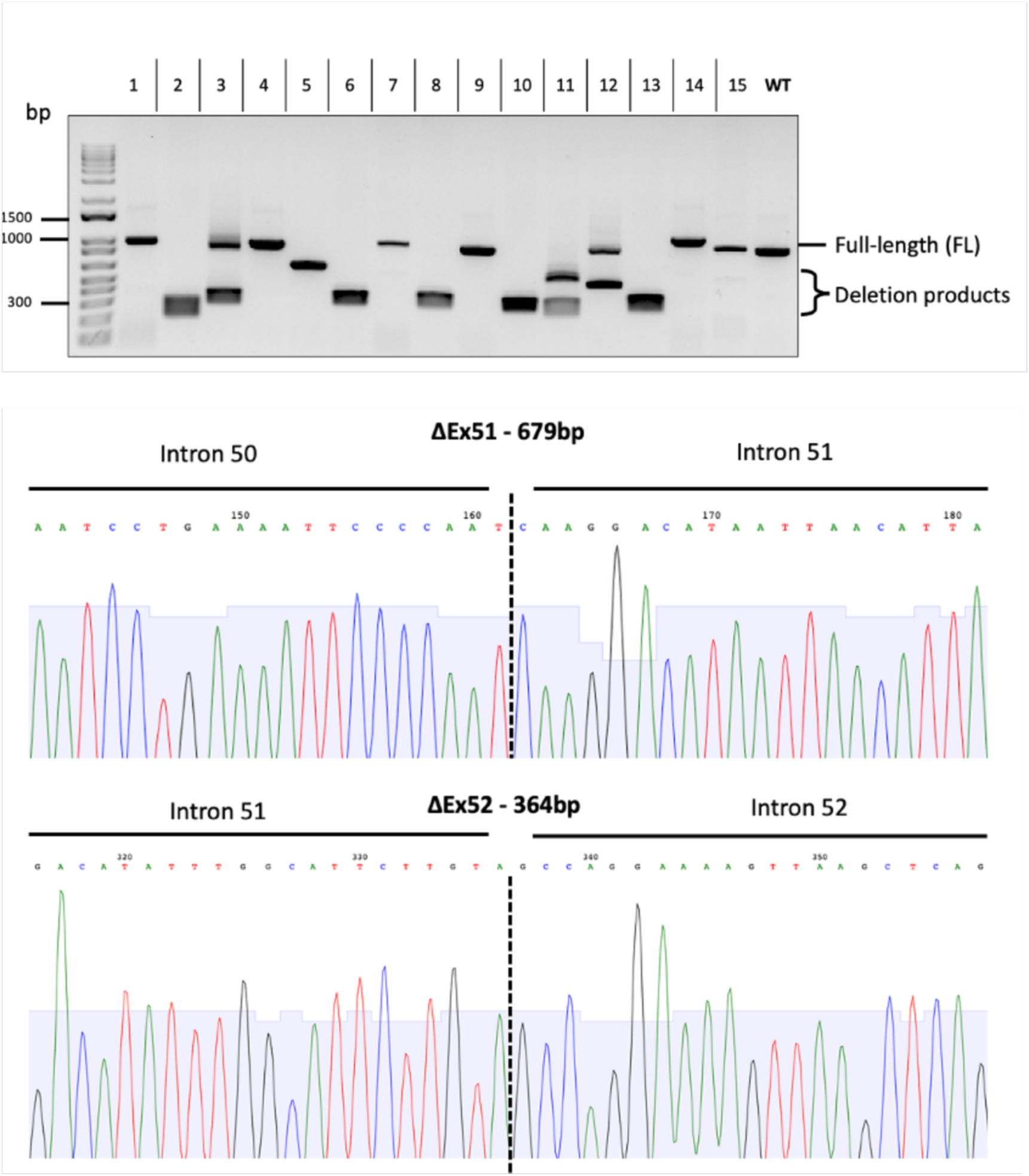
Genotyping of the founders by PCR. **(A)** PCR primers spanned the cut sites to capture the intervening deletion resulting from the dual cut. The last sample represents the control WT band of 965bp. (B) Sanger sequencing further confirmed the absence of exon 51 (679bp deletion) and exon 52 (364bp deletion) from the genomic sequence.

**Table 2.**
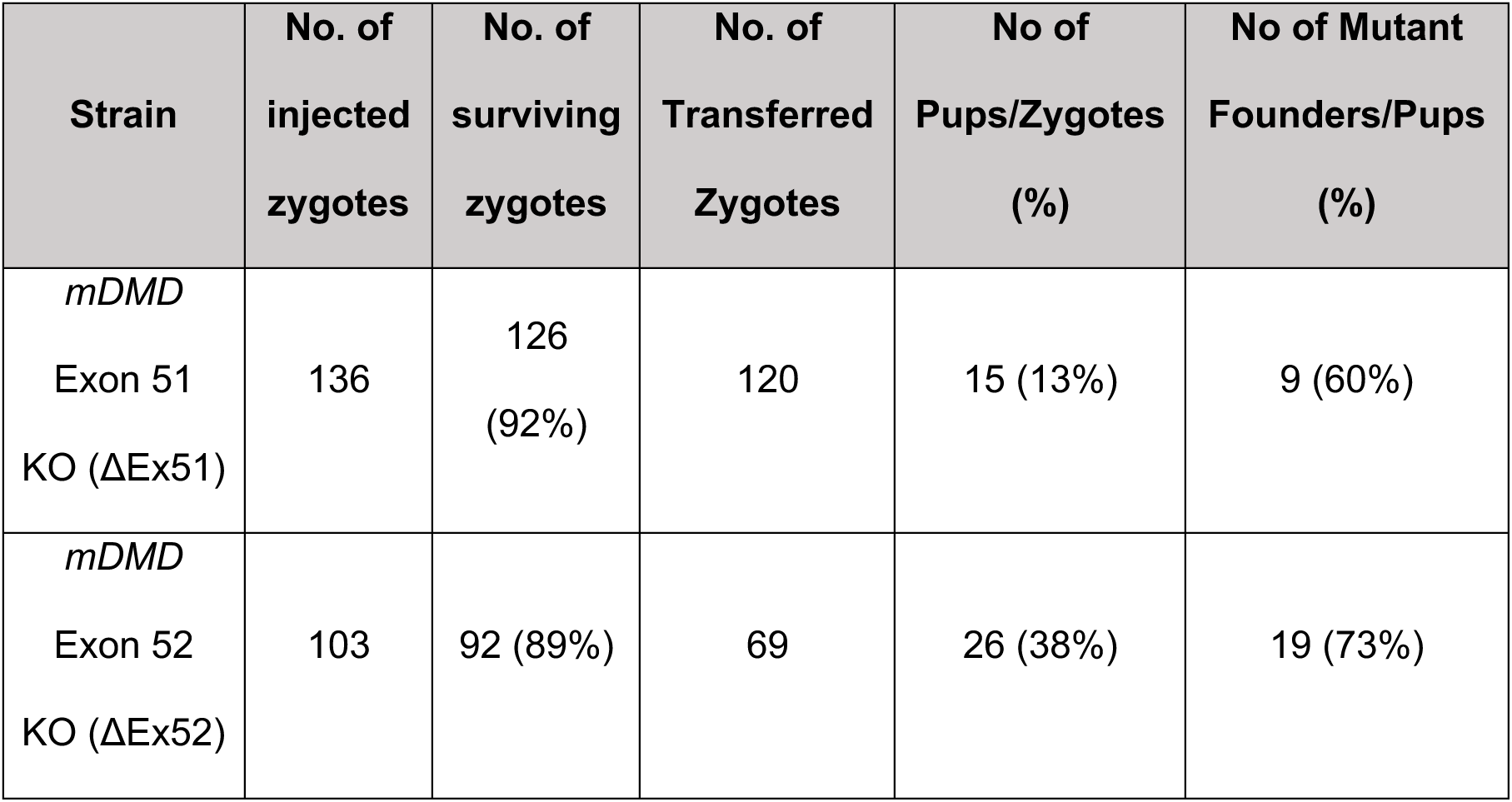
Efficiency of CRISPR-Cas9 mediated genomic editing by cytoplasm microinjection.

| Strain | No. of injected zygotes | No. of surviving zygotes | No. of Transferred Zygotes | No of Pups/Zygotes (%) | No of Mutant Founders/Pups (%) |
| --- | --- | --- | --- | --- | --- |
| *mDMD*<br>Exon 51<br>KO (ΔEx51) | 136 | 126<br>(92%) | 120 | 15 (13%) | 9 (60%) |
| *mDMD*<br>Exon 52<br>KO (ΔEx52) | 103 | 92 (89%) | 69 | 26 (38%) | 19 (73%) |

### Molecular characterisation of the generated DMD mouse models

To confirm the downstream molecular consequence of the genomic exon removal on dystrophin expression, we performed rigorous molecular analyses on muscle tissues such as quadriceps, tibialis anterior, triceps and hearts. RT-PCR across the target region demonstrated a reduction in amplicon size consistent with the absence of the deleted exons in the *Dmd* transcript (Fig 3A, 3B). Sanger sequencing of the RT-PCR products corroborated these findings, confirming the expected exon deletion on RNA level and the joining of the neighbouring exons (data not shown). Notably, RT-qPCR across the terminal end of the transcript revealed a substantial reduction (85%) in the 3’ end of the *Dmd* transcript, (Fig 3C). Subsequent protein-level characterisation via Western blotting and immunohistochemistry validated the absence of dystrophin in tissues from exon-51 or exon-52 deletion mouse models (Figs 4 and 6). These analyses confirmed the successful knockout and absence of dystrophin in the generated models.

**Figure 3:**
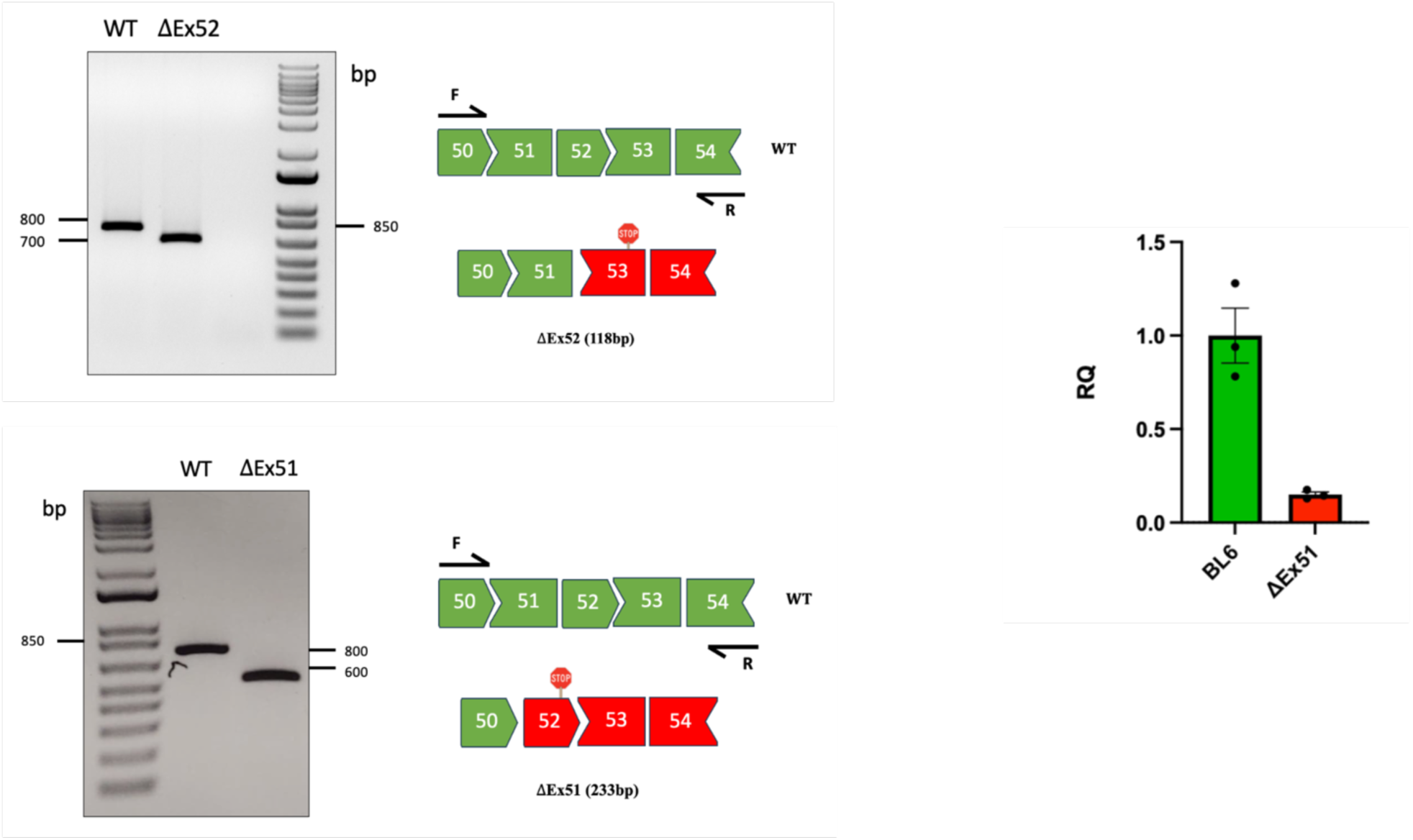
RT-PCR analysis to validate the deletion of exons 51 and 52. (A, B) RT-PCR primers were in exons 50 (F) and 54 (R), and the amplicon size is 827 bp for WT mice and 594 bp for ΔEx51 DMD mice, and 709bp for ΔEx52 DMD mice. RT-PCR products are schematised next to each gel image. **(C)** RT-qPCR analysis of transcript level differences between WT and KO, relative to the endogenous control Beta-actin. Primers span the terminal end of the DMD gene in exons 77 and 78.

**Figure 4:**
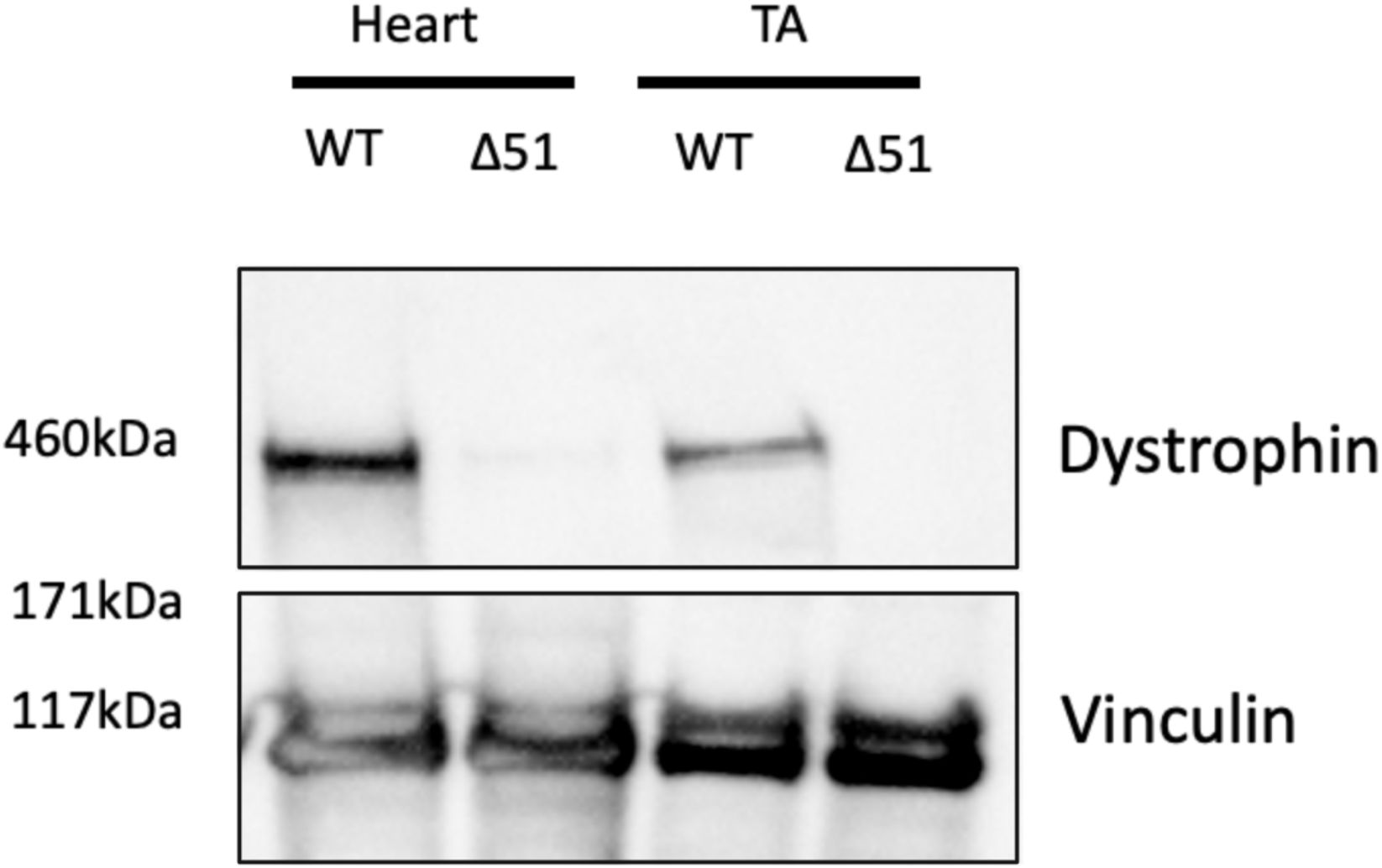
Western blot from protein lysates of ΔEx51 mice and healthy controls. 50ug of total protein was loaded per lane. There is absence of dystrophin (427kDa) in the tibialis anterior (TA) muscle and cardiac tissues compared to controls. Vinculin was used as the loading control for normalisation.

### Phenotypic characterisation of the generated DMD mouse models

It is known that mouse models lacking dystrophin, particularly those with a C57BL/6 background, do not exhibit severe phenotypes of muscle weaknesses and premature death observed in human DMD patients (8, 10). However, dystrophin-deficient mice display measurable characteristics akin to DMD, including weaker grip strength, elevated serum CK levels, and abnormal muscle histology (10). We systematically evaluated these parameters in our knockout mouse models and observed similar findings. Normalised grip strength assessment revealed a significant reduction in 4-week-old knockout mice compared to their WT counterparts with an average difference of 2-fold in grip strength and 2.5-fold increase in peak force generated across each of the three pulls, indicating compromised muscle performance due to dystrophin deficiency (Fig 7). Additionally, serum CK levels collected from tail bleeds of the 4-week-old mice confirmed elevated levels in knockout mice of up to 16-fold compared to the WT controls (Fig 8). Histological analysis across muscle tissue sections of our mutant models consistently revealed dystrophic pathology characterised by centrally nucleated myofibers, fibrotic deposition, and variable myofiber size (Fig 5). Furthermore, immunofluorescence labelling showed a lack of dystrophin in the mutant models of DMD compared with defined borders present across the sarcolemma in healthy muscle (Fig 6). These comprehensive characterisations collectively affirm the dystrophic phenotypes evident in our knockout models.

**Figure 5:**
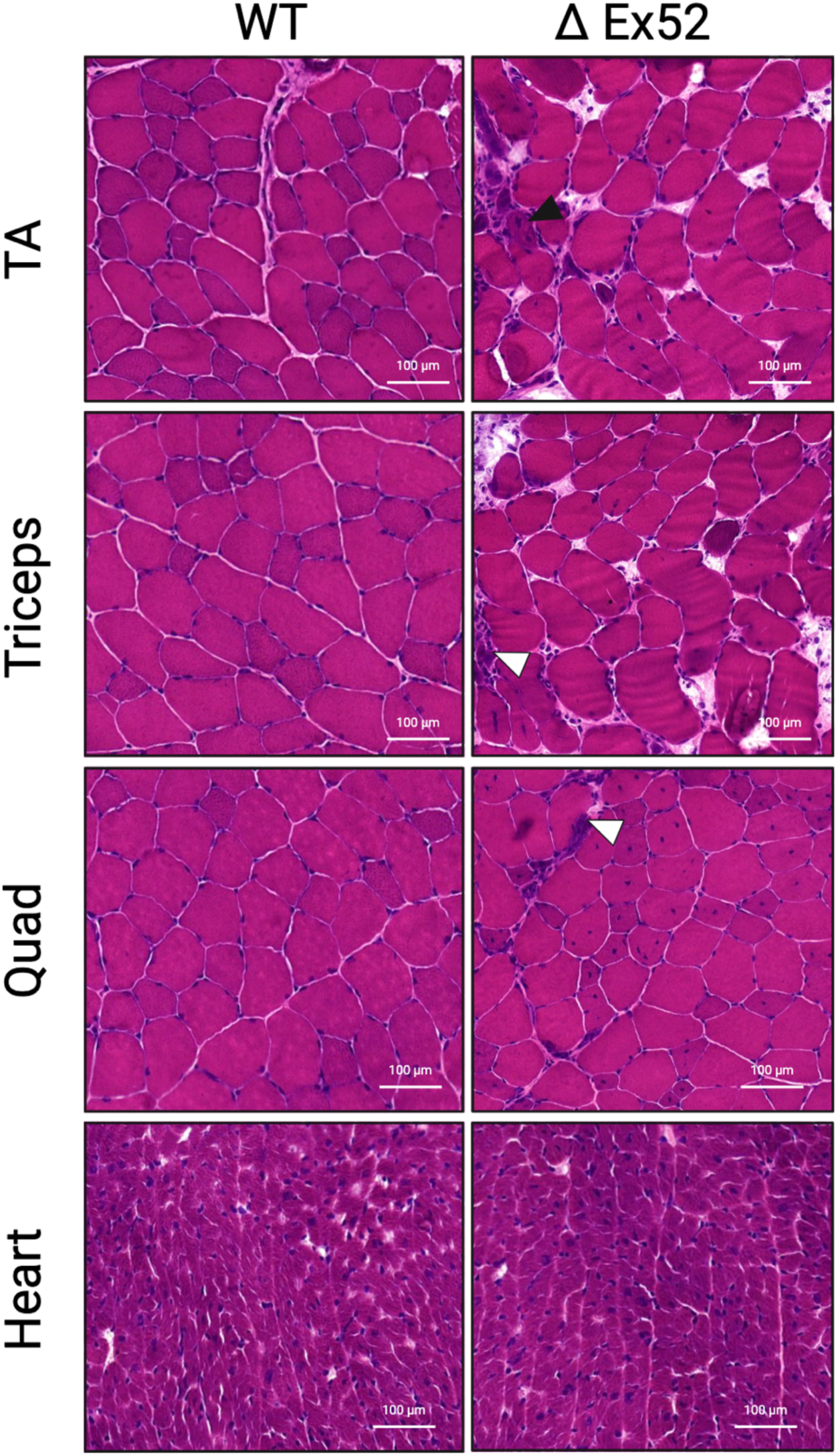
Hematoxylin and eosin (H&E) stained transverse cryosections of whole tibialis anterior (TA), triceps, heart and quadriceps of WT and ΔEx52 DMD mice. Note extensive inflammatory infiltrate, increased necrosis and fibrosis, and centralised myonuclei in DMD mice.

**Figure 6:**
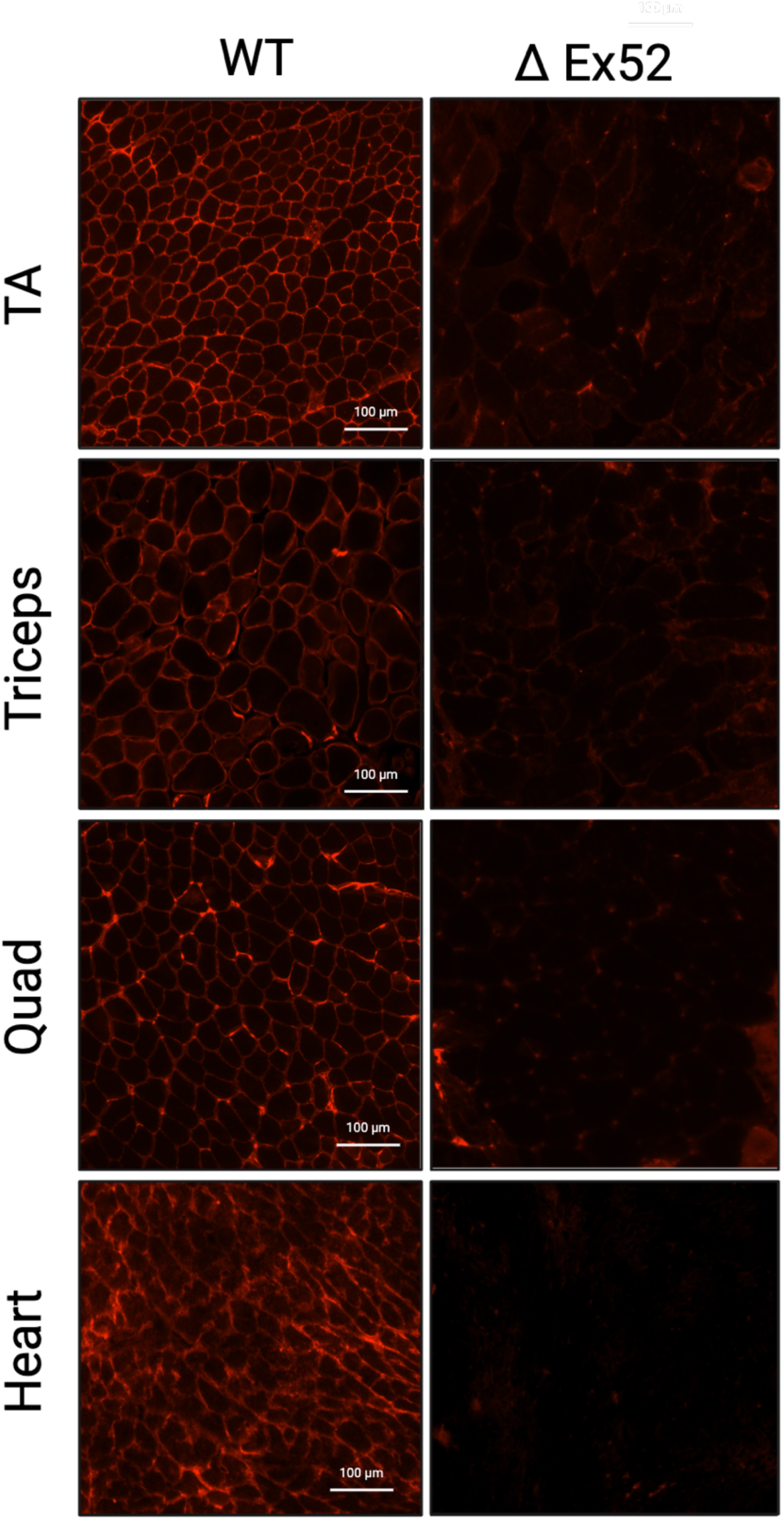
Immunofluorescence indicates the absence of dystrophin in muscle tissues of ΔEx52 DMD mice. Samples were snap-frozen in isopentane suspended within liquid nitrogen. Dystrophin is shown in red: scale bar, 100 μm.

**Figure 7:**
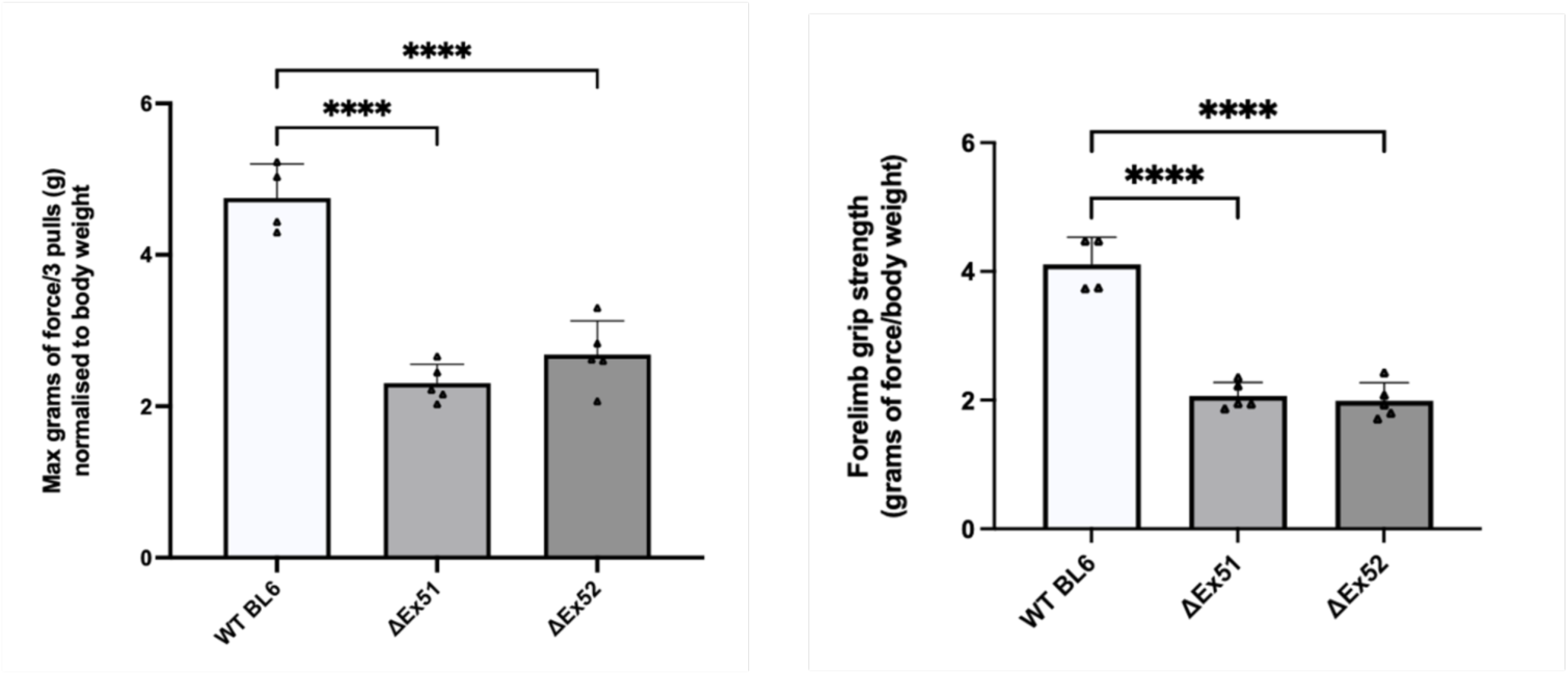
Forelimb grip strength testing normalised to body weights. **(A)** Maximum forelimb pulls from 3 sets (4 pulls/set) each across WT, ΔEx51 and ΔEx52 mice. **(B)** Forelimb grip strength analysis. Data are represented as mean ± SEM. Unpaired Student’s *t*- test was performed. \*\*\**P* < 0.05 (*n* = 5).

**Figure 8:**
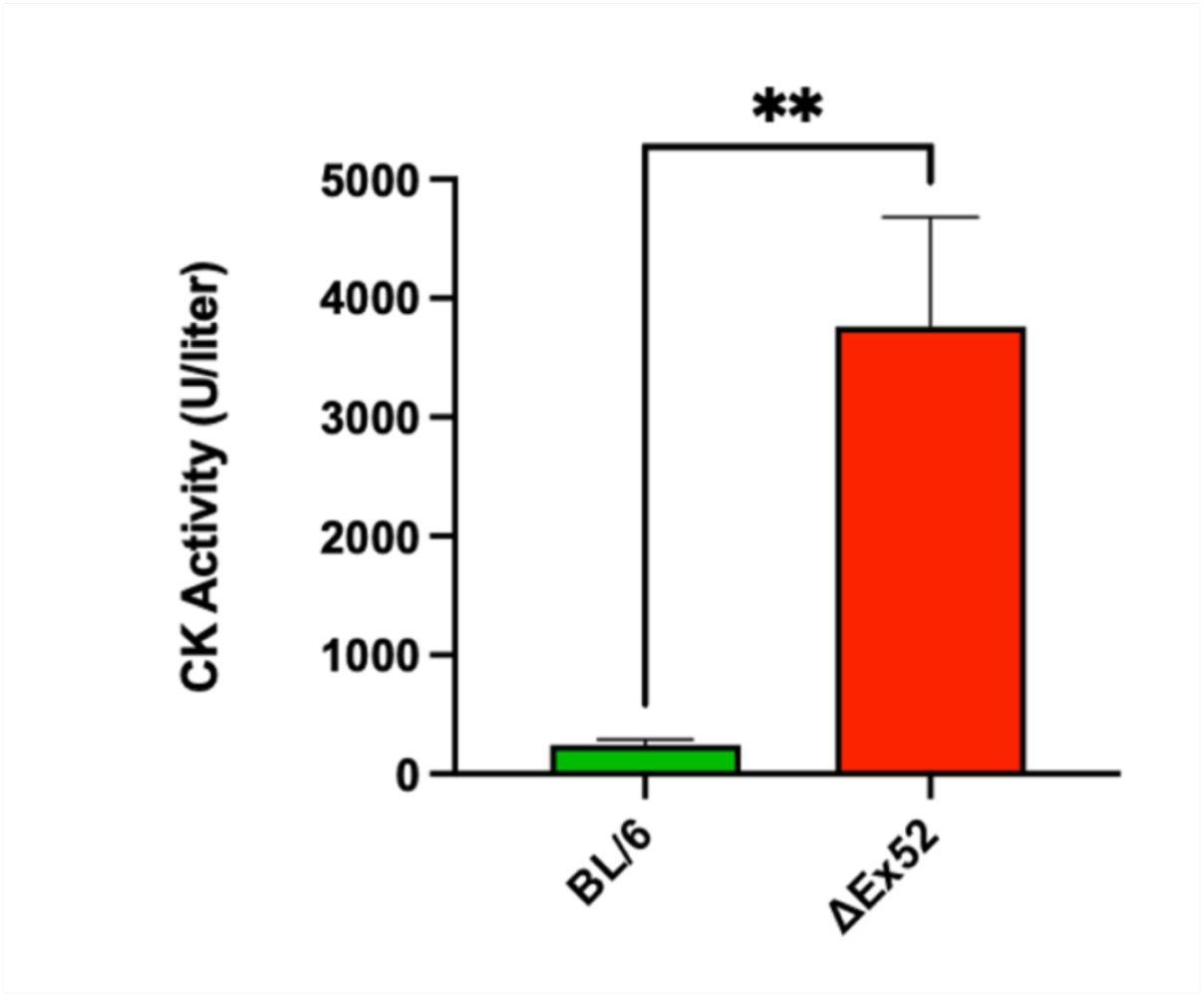
Serum creatine kinase (CK), a marker of muscle damage and membrane leakage, was measured in WT (C57BL/6), ΔEx51, and ΔEx52 mice. Data represented as mean ± SEM. Unpaired Student’s *t*-test was performed. \**P* < 0.005 (*n* = 5)

## Discussion

In this study we generated mouse models of DMD lacking exon 51 and exon 52 of the murine dystrophin gene, representing one of the most prevalent hotspot regions of exonic deletion mutations in DMD patients. These ΔEx51 and ΔEx52 DMD male mice display the hallmarks of DMD, where the severity of the disease, as marked by the absence of dystrophin protein expression, muscle histology and serum CK, were comparable to other published murine models of DMD (15, 17, 27). In a therapeutic context, correcting exon 51 and exon 52 deletions through exon skipping or reframing surrounding exons could potentially treat 8% and 12% of patients with DMD, respectively (18). However, it is essential to recognise that these mouse models are not the most effective pre-clinical models when testing sequence-dependent therapies, as mouse and human DMD sequences are not identical. As such, mouse models harbouring targetable human DMD sequences are an attractive platform for testing genome editing strategies and other gene therapies in a more clinically relevant context (10, 16). This protocol can also be applied to generate and characterise such models.

Identifying hemizygous males, who carry a single copy of the deletion, and homozygous females, who harbour the deletion on both X chromosomes, is vital for breeding purposes to maintain the DMD deletion line for further study. Discerning these specific genotypes within the founder animals enables controlled breeding to perpetuate the deletion line in subsequent generations. However, it is important to note that CRISPR-Cas9 cutting frequently induces unintended on-target large deletions that could span kilo/mega bases that will remove the primer binding site which will be undetected by our standard PCR (19).

We observed significantly reduced transcript levels in our DMD mouse models, presumably due to Nonsense-Mediated Decay (NMD) (20). This assay may be a valuable tool for quantifying the amount of transcript restoration when testing therapeutic guide candidates.

However, a recent investigation into PTC-containing transcripts revealed that inhibiting NMD did not normalize DMD expression in an *mdx* mouse model. Instead, Spitali and colleagues (2020) hypothesised that the transcript reduction occurs through an NMD-independent epigenetically mediated mechanism through histone methylation (21).

Dystrophin typically exhibits a molecular weight of 427 kDa, corresponding to the major Dp427 muscle protein isoform. In muscle samples from individuals with Duchenne muscular dystrophy (DMD), the average dystrophin levels are approximately 1.3%, ranging from 0.7% to 7% of the average observed in healthy muscle (22). Notably, dystrophin expression is a frequently employed secondary outcome measure in various clinical trials and in vivo therapeutic assessments (23). Assessing the intensity of the dystrophin band on a blot should provide crucial information on the presence or absence of full-length dystrophin, where non-muscle dystrophin isoforms may also be expressed. Our DMD models show an absence of full-length dystrophin in both heart and TA samples. Recently, the emergence of the capillary western immunoassay has offered high-throughput analysis of dystrophin quantitation, offering greater sensitivity in detecting low-abundance proteins. This platform needs 100-fold less sample and 500-fold less antibody to detect dystrophin (22). This automated system may provide a more accurate assessment of dystrophin expression, especially that of low-level dystrophin restoration, when assessing multiple therapeutic interventions.

Proper processing of the muscle tissue is important to study muscle biology and pathology of DMD. In alignment with what others have seen, we show that rapid freezing of freshly isolated skeletal muscle tissue using isopentane pre-cooled with liquid nitrogen and tragacanth gum was optimal for preserving tissue integrity (24). Immersing the muscle directly into isopentane ensured rapid freezing, preventing the formation of ice crystals and ruptured cell membranes (30). In the context of our IF results, all muscle tissues from the mutant models lack subcellular protein localisation within the sarcolemma. Haematoxylin and Eosin (H&E) staining is the gold standard approach to investigate general dystrophic pathology, giving vital information on the location of the nucleus within the fiber, muscle fiber size and fibrosis deposition (3). In our mutant model, these hallmarks included centrally nucleated myofibers, fibrosis deposition and variable myofiber size.

The most common biomarker in the clinical diagnosis of DMD is the serum activity levels of creatine kinase. In short, muscle contraction without dystrophin tears the sarcolemma to release creatine kinase into the bloodstream (1, 3). Across other publications outlining other dystrophic models of DMD, we see a stark variation in CK values ranging from as low as 8 to a 40-fold difference compared to WT mice (26, 27). Here we sampled an alternative route of blood collection through the lateral tail vein. The standard collection route, as recommended by TREAT-NMD has been cardiac puncture or used by labs, including submandibular vein collection. The latter method requires an anaesthesia system, but both have the limitation of being unsuitable for frequent small blood volume collection when wanting to assess CK levels at multiple time points. Our approach offers an alternative that is not limited to endpoint analysis. As evidenced in other dystrophic murine models, the analysis of serum CK levels is expected to reveal a significant elevation compared to WT mice, indicating muscle damage and membrane instability.

Much like individuals affected by Duchenne Muscular Dystrophy (DMD), the absence of dystrophin in the muscle fibres of these DMD mice renders them susceptible to exercise-induced damage, resulting in impaired muscle function compared to WT mice (3, 10). A non-invasive functional measure such as forelimb grip strength can be used assess this impairment and monitor the disease’s natural progression without influencing muscle pathology. Our mutant models demonstrated a noticeable reduction in the KO models compared to WT mice. This comprehensive phenotypic characterisation provides valuable insights into DMD pathology and is a pivotal tool to actively gauge the efficacy of specific treatments, including CRISPR therapy and oligonucleotide-skipping therapy. Utilising this protocol, one can actively measure the extent of therapeutic rescue, offering a dynamic assessment of treatment outcomes and advancing our understanding of potential interventions for further development. Nevertheless, additional efforts are needed to fully characterise the DMD pathology in these models. These include a long term follow up of dystrophic mice, cardiac pathology assessment and evaluation of muscle physiology.

Off-targets are an important consideration when generating bespoke cell/animal models using CRISPR-Cas9 because the observed phenotypes from genome modifications could potentially stem from off-target effects rather than on-target modifications (31). Therefore, it is crucial to minimise the off-target activity by performing extensive in-silico screening using reliable bioinformatic tools to identify gRNAs with limited potential off-target binding sites. In the context of generating a mutant mouse model, off-target effects at other genomic loci can be mitigated by successive backcrossing in the breeding process to segregate away off-target mutations that may have occurred during founder generation.

## Declarations

Ethics approval: Animal work described in this manuscript has been approved and conducted under the oversight of the Animal Ethics Committee of South Australian Health and Medical Research Institute (SAHMRI) and The University of Adelaide.

Consent to participate: Not applicable

Consent for publication: Not applicable

## Acknowledgement

We would like to thank SAHMRI for providing the facilities to carry out leading biomedicals research. We would also like to thank the South Australian Genome Editing Facility (SAGE) for assistance with generating these gene-edited models. We would like to thank Annemieke Aartsma-Rus for providing useful comments on the manuscript. BioRender was used to generate figures for this manuscript.

## Competing interests

The authors have declared that no competing interests exist.

Availability of data and materials: The datasets supporting the conclusions of this article are included within the article and its supporting information files.

Authors’ contributions

JA Contributed to conceptualisation, contributed significantly to methodology, performed majority of investigation, formal data analysis, and data interpretation, handled all data curation, wrote first draft of the manuscript, and created all visualisations. JA also reviewed all other author manuscript suggestions and edited the final draft. YCJC contributed to methodology and reviewed manuscript during drafting. SGP performed investigation on some aspects – including performing all microinjections. PQT and FA performed the majority of conceptualisation and methodology, project administration and supervision as well as contributed to data interpretation. Both PQT and FA performed significant revision and editing to manuscript during drafting. All authors read and approved the final manuscript.

## List of abbreviations

DMD: Duchenne Muscular Dystrophy
CRISPR: Clustered regularly interspaced short palindromic repeats
sgRNA: Single guide RNA
DSBs: Double stranded breaks
NHEJ: Non-homologous end joining
InDels: Insertions/Deletions
IVT: In vitro-transcription
HRP: Horseradish Peroxidase
TA: Tibialis anterior
NMD: Nonsense-mediated decay
PTC: Premature termination codon
PCR: Polymerase chain reaction
RT-PCR: Reverse transcription polymerase chain reaction
qRT-PCR: Quantitative reverse transcription polymerase chain reaction
PFA: Paraformaldehyde
H&E: Haematoxylin and Eosin
WT: wildtype
LN: Liquid Nitrogen
RFLP: Restriction fragment length polymorphism

## Supporting information

The protocols in PDF format available from protocols.io are provided as Supporting Information file 1.

**Supporting Information file 1:** Step-by-step protocols, also available on protocols.io

**S1 Table:**
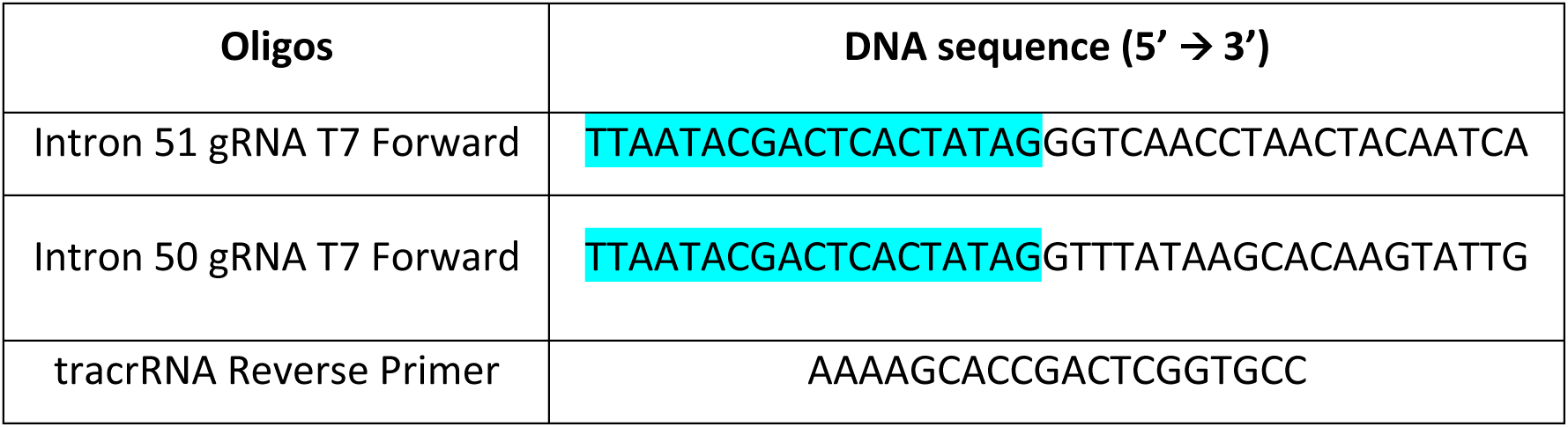
Mouse Zygote injection primers and guide oligos to generate ΔEx51 DMD mice. The T7 primer sequence is highlighted in blue.

**S2 Table:**
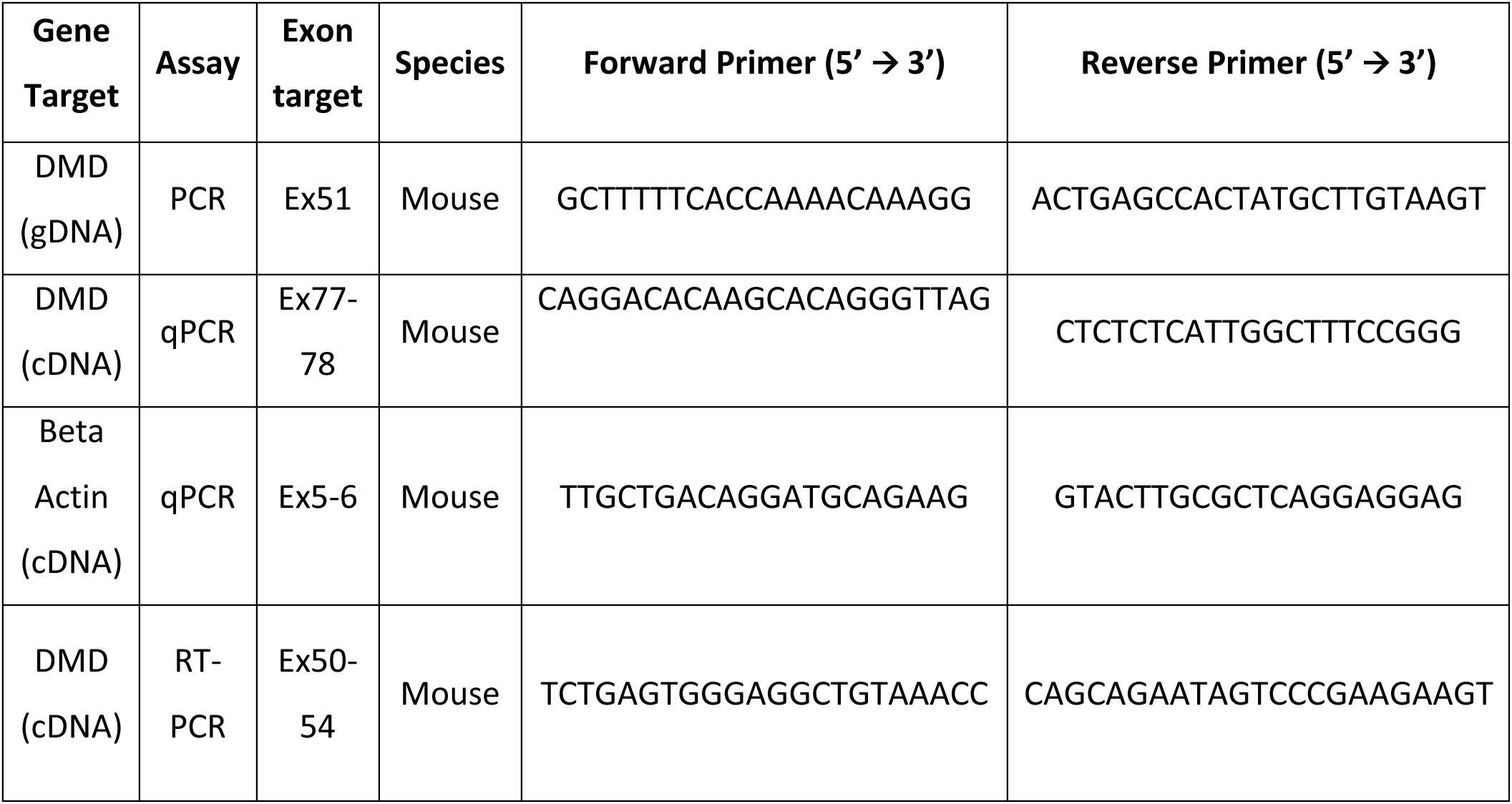
PCR, RT-PCR primers and SYBR Green RT-qPCR Primers.

**S3 Table:** Antibodies used for Immunofluorescence and Western Bot analyses.

| **Immuno Antibody** | **Species** | **Company** | **Catalog #** | **Dilution Factor** |
| --- | --- | --- | --- | --- |
| MANDYS8 | Mouse | Sigma-Aldrich | D8168 | 1:100 |
| Donkey anti-Mouse IgG (H+L)<br>Alexa Fluor™ 594 | Donkey | Thermo Fisher Scientific | R37115 | 1:300 |
| **Western Antibody** | **Species** | **Company** | **Catalog #** | **Dilution Factor** |
| MANDYS8 | Mouse | Sigma-Aldrich | D8168 | 1:1000 |
| Goat anti-Mouse IgG (H+L) HRP | Donkey | Thermo Fisher Scientific | 31430 | 1:8000 |

