## Supporting Information File 1 for "Generation and characterisation of mouse models of Duchenne Muscular Dystrophy (DMD)"

### In vitro Transcription (IVT) of Guide RNAs for Cytoplasmic Microinjection

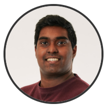

Jayshen Arudkumar
University of Adelaide, South Australian Health and Medical ...

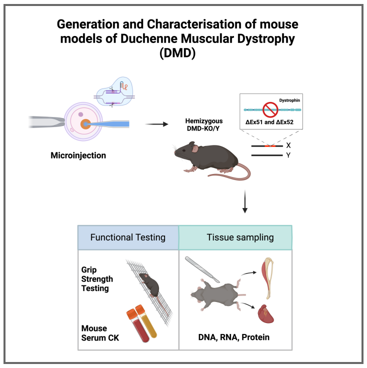


**Protocol Info:** Jayshen Arudkumar, Yu Chinn Joshua Chey, Sandra Piltz, Paul Quinton Thomas, Fatwa Adikusuma . In vitro Transcription (IVT) of Guide RNAs for Cytoplasmic Microinjection. **protocols.io** https://protocols.io/view/in-vitro-transcription-ivt-of-guide-rnas-for-cytop-c7jnzkme

**Created:** Jan 15, 2024

**Last Modified:** Jan 29, 2024

**PROTOCOL integer ID:** 93518

**Keywords:** CRISPR, Microinjection, mouse, embryo, DMD, Phenotyping, Knockout

#### DISCLAIMER

These protocols are for research purposes only.

#### ABSTRACT

**Paper abstract:** CRISPR-Cas9 gene-editing technology has revolutionised the creation of precise and permanent modifications to DNA, enabling the generation of diverse animal models for investigating potential treatments. Here, we provide a protocol for the use of CRISPR-Cas9 to create murine models of Duchenne Muscular Dystrophy (DMD) along with a step-by-step guide for their phenotypic and molecular characterisation. The experimental procedures include CRISPR microinjection of embryos, molecular testing at the DNA, RNA, and protein levels, forelimb grip strength testing, immunostaining and serum creatine kinase (CK) testing. We further provide suggestions for analysis and interpretation of the generated data, as well as the limitations of our approach. These protocols are designed for researchers who intend on generating and using mouse models to study DMD as well as those seeking a detailed framework of phenotyping to contribute to the broader landscape of genetic disorder investigations.
**Protocol summary:** In this section we present a protocol for CRISPR injection into mouse zygotes to create the ΔEx51 DMD model. Here, we successfully generated guide RNAs (sgRNAs) targeting intronic regions flanking the mouse Dmd target region of exon 51 to create a large intervening deletion.

#### IMAGE ATTRIBUTION

BioRender was used to generate figures for this manuscript.

MATERIALS

1. NEB HiScribe™ T7 Quick High Yield RNA Synthesis Kit
2. Qiagen RNEasy Mini Kit RNA Cleanup (Qiagen)
3. Purified pX459V2 construct (backbone from Addgene Plasmid #62988)
4. T7 primer + FWD and REV sequences with overhangs
5. NEB Formaldehyde Load Dye
6. QIAquick Gel Extraction Kit (Qiagen)
7. QIAQuick PCR Purification Kit (Qiagen)

SAFETY WARNINGS

Wear proper PPE (gloves, safety goggles, enclosed shoes and lab coat) and prepare solvents in a chemical fume hood. Dispose used solvents or waste material in an appropriate biohazard waste containers.

ETHICS STATEMENT

Animal work described in this manuscript has been approved and conducted under the oversight of the Animal Ethics Committee of South Australian Health and Medical Research Institute (SAHMRI) and The University of Adelaide.

#### Choosing a sgRNA pairing to create the deletion

1 The in-silico design of guide RNAs was conducted through various online guide design tools and narrowed down based on their predicted on-target and off-target activities. Tools that we would recommend for this task include Benchling and CRISPOR (1). The phospho-annealed guide oligos were then cloned into BbsI-linearised pX459V2 plasmid using the Golden Gate Assembly method. Note that ‘CACC’ and ‘AAAC’ overhangs allow the oligos to bind the complementary overhanging DNA at the cut sites in the plasmid created by the BbsI digestion (2). This process yielded two recombinant constructs, each housing the left intron and right intron, respectively.
gRNA 1 (Right Intron)
Position is 156bp away from 3' exon 51
Forward: 5' - GGTCAACCTAACTACAATCA - 3'
Rev comp: 5' - TGATTGTAGTTAGGTTGACC - 3'

Forward oligo: 5' -CACCGGTCAACCTAACTACAATCA - 3'
Reverse oligo: 5' -AAACTGATTGTAGTTAGGTTGACC - 3'

gRNA 2 (Left Intron)
Position is 290bp away from 5' exon 51
Forward: 5' - GTTTATAAGCACAAGTATTG - 3'

Rev comp: 5' - CAATACTTGTGCTTATAAAC - 3'

Forward oligo: 5' - CACCGTTTATAAGCACAAGTATTG - 3'
Reverse oligo: 5' - AAACCAATACTTGTGCTTATAAAC - 3'

#### PCR amplification of guide template

1

**Note**

The oligos used for Forward and Reverse PCRs are seen in Supplementary Table S1. Here is an expanded form:

gRNA 1 (Right Intron)
T7 Guide Primer for Forward PCR:
5' - TTAATACGACTCACTATAGGGTCAACCTAACTACAATCA - 3'

gRNA 2 (Left Intron)
T7 Guide Primer for Forward PCR:
5' - TTAATACGACTCACTATAGGTTTATAAGCACAAGTATTG - 3'

tracrRNA Reverse Primer: 5' - AAAAGCACCGACTCGGTGCC - 3'

2 Prepare PCR mix using the T7 Guide Primer for Forward PCR and tracrRNA reverse primer. The T7 promoter is introduced into the template. Add 1 µL of each of the miniprepped recombinant guide plasmids (at ~1-3 ng/µL) to 1 tube each and 1 µL Ultrapure water to the final tube.

| A | B |
|---|---|
| Reagent | Amount |
| Ultrapure water | 12.3 µL |
| NEB Phusion HF Reaction Buffer (5x) | 4 µL |
| T7 guide primer (10 µM) | 1 µL |
| tracrRNA reverse primer (10 µM) | 1 µL |
| Roche PCR Grade Nucleotide Mix (10 mM) | 0.5 µL |
| NEB Phusion HF DNA Polymerase (2 U/µL) | 0.2 µL |
| Total | 19 µL |

3 Place the tubes in a thermocycler with the following parameters:

| A | B | C |
|---|---|---|
| Steps | T7 PHUSION | |
| 1 | 98°C | 3 min |

| A | B | C |
|---|---|---|
| 2 | 98°C | 15 s |
| 3 | 60°C | 20 s |
| 4 | 72°C | 15 s |
| 5 | Go to step 2 | 32 times |
| 6 | 72°C | 5 min |
| 7 | 4°C | ∞ |

#### PCR amplification of guide template

4 Make a 1% agarose gel and run 5 µL of the PCR products 40 min @100 V.

**Note**

Testing to see if the plasmid has the correct insert. Band should be present at approximately 100 bp

5 Combine the remainder of all PCR reactions with the correct band and perform Qiagen PCR Purification on the mixture. Use NanoDrop to measure concentration of DNA. A260/A280 values should fall in the range of 1.8-2.0.

**Note**

This is to confirm if the DNA is still present and determine the amount needed for the IVT reaction.

#### In vitro transcription (IVT)

6 Transfer 2 µL to a PCR tube for testing later.

**Note**

We are using the purified PCR product as the template for IVT with the NEB HiScribe™ T7 Quick High Yield RNA Synthesis Kit.

7 Mix the following reagents in a PCR tube:

| A | B |
|---|---|
| Reagent | Amount |
| Ultrapure H2O | Up to 60 µL |
| IVT gRNA Product | 40 µL |
| NEB DNase I (RNAse-free) (2 U/µL) | 4 µL |
| Total | 104 µL |

Note

Degrades DNA

8 Incubate 15 min @ 37 °C.

9 Transfer 2 µL to a separate PCR tube for testing later.

10 Perform Qiagen RNEasy Mini Kit RNA Cleanup, eluting in 30 µL water.

11 Transfer 2 µL to a separate PCR tube for testing later.

12 Add 2 µL NEB Formaldehyde Load Dye to each of the 2 µL RNA aliquots set aside.

13 Place the mixture in a thermocycler with the following parameters:

| Steps | Reagent | Amount |
|---|---|---|
| 1 | 85°C | 3 min |
| 2 | 4°C | ∞ |

14 Make an RNase-free 1% agarose gel and run RNA aliquots 30 min @ 100 V.

Note

Testing the gRNA has been produced correctly. Bands should be present at >100 bp (RNA will run higher on the gel than DNA when run against a DNA ladder).

#### Cytoplasmic Injection

15 On the day of zygote injection, prepare the injection mix as follows:

| A | B |
|---|---|
| Reagent | Amount |
| Ultrapure H2O | 10.125 µL |
| Injection buffer (10x) | 1.5 µL |
| gRNA 1 | To give 50 ng/µL |
| gRNA 2 | To give 50 ng/µL |
| SpCas9 mRNA | To give 100 ng/µL |

| A | B |
| --- | --- |
| Total | 15 µL |

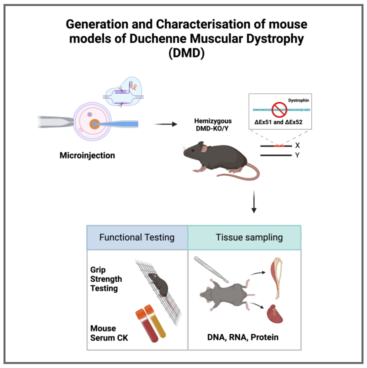


### Genomic DNA Extraction and Genotyping PCR on Mouse Tissues

Jayshen Arudkumar[1,2], Yu Chinn Joshua Chey[1,2], Sandra Piltz[1,2], Paul Quinton Thomas[1,2], Fatwa Adikusuma[1,2]

[1]University of Adelaide; [2]SAHMRI

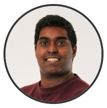

Jayshen Arudkumar
University of Adelaide, South Australian Health and Medical ...

**Protocol Info:** Jayshen Arudkumar, Yu Chinn Joshua Chey, Sandra Piltz, Paul Quinton Thomas, Fatwa Adikusuma . Genomic DNA Extraction and Genotyping PCR on Mouse Tissues. **protocols.io** https://protocols.io/view/genomic-dna-extraction-and-genotyping-pcr-on-mouse-c8d9zs96

**Created:** Jan 30, 2024

**Last Modified:** Jan 30, 2024

**PROTOCOL integer ID:** 94369

**Keywords:** Genomic, CRISPR, Mouse, DNA, Therapy, DMD

#### DISCLAIMER

These protocols are for research purposes only.

#### ABSTRACT

**Paper abstract:** CRISPR-Cas9 gene-editing technology has revolutionised the creation of precise and permanent modifications to DNA, enabling the generation of diverse animal models for investigating potential treatments. Here, we provide a protocol for the use of CRISPR-Cas9 to create murine models of Duchenne Muscular Dystrophy (DMD) along with a step-by-step guide for their phenotypic and molecular characterisation. The experimental procedures include CRISPR microinjection of embryos, molecular testing at the DNA, RNA, and protein levels, forelimb grip strength testing, immunostaining and serum creatine kinase (CK) testing. We further provide suggestions for analysis and interpretation of the generated data, as well as the limitations of our approach. These protocols are designed for researchers who intend on generating and using mouse models to study DMD as well as those seeking a detailed framework of phenotyping to contribute to the broader landscape of genetic disorder investigations.

#### IMAGE ATTRIBUTION

Image was generated using Biorender.

#### GUIDELINES

This protocol is designed for use with 25-50 mg of ear or tail tissue. The High Pure PCR Template Preparation kit (Roche) was followed for DNA purification. PCR conditions may vary based on considerations such as primer design and the type of polymerase employed in amplification.

MATERIALS

- PPE (Personal Protective Equipment)
- High Pure PCR Template Preparation Kit (Roche)

SAFETY WARNINGS

Wear proper PPE (gloves, safety goggles, enclosed shoes and lab coat) and prepare solvents in a chemical fume hood. Dispose used solvents or waste material in an appropriate biohazard waste containers.

ETHICS STATEMENT

Animal work described in this manuscript has been approved and conducted under the oversight of the Animal Ethics Committee of South Australian Health and Medical Research Institute (SAHMRI) and The University of Adelaide.

#### Lysis and Protein Digestion

1 Add 200 µL of Roche Tissue Lysis Buffer to a 1.5 mL tube containing the tissue sample.

Note

Ensure the buffer maintains a pH of 7.4, as Tris-HCl buffers the pH Note: EDTA chelates Mg2+/Ca2+ (essential for DNases), and urea lyses cells while disrupting hydrophobic interactions to destabilise proteins, including nucleases.

Note

For a higher DNA yield, it's advisable to chop up the tissue sample.

2 Add 40 µL of approximately 413 µg/µL Roche Proteinase K solution.

3 Vortex the tube to mix the contents.

4 Incubate the tube at 55 °C for 3 hours.

Note

Longer incubation times, including overnight or multiple days, can increase DNA yield as Proteinase K degrades proteins, solubilising them (including DNases).

5 Incubate a sufficient amount of Roche Elution Buffer in a 1.5/10 mL tube at 75 °C until ready for use.

6 Vortex the tube

#### DNA Column Binding

7 Add 200 µL of Roche Binding Buffer to the lysed sample.

**Note**

Ensure a pH of 4.4 is maintained with Tris-HCl

**Note**

Guanidinium chloride and urea disrupt hydrophobic interactions, destabilizing proteins and disrupting hydrogen bonds between DNA and water, allowing stronger binding to the silica filter. Triton-X 100 solubilizes lipids, allowing them to flow through the filter tube.

8 Add 100 µL of isopropanol.

9 Vortex the tube

10 Centrifuge the tube at 13,000 RCF for 5 minutes.

11 Decant the solution into a Roche High Pure Filter Tube in a collection tube.

12 Centrifuge the tube at 8,000 RCF for 1 minute.

**Note**

Soluble contaminants, including proteins, lipids, and polysaccharides, are washed through the filter, while protein and salt residues remain with the DNA.

13 Discard the flow-through and place the filter tube in a new collection tube.

14 Add 500 µL of Roche Inhibitor Removal Buffer to the filter tube.

**Note**

Guanidium chloride removes residual proteins and pigments. Ethanol removes the salts.

15 Centrifuge the tube at 8,000 RCF for 1 minute.

16 Discard the flow-through and place the filter tube in a new collection tube.

#### Additional Washing

17 Add 500 µL of Roche Washing Buffer to the filter tube.

18 Discard the flow-through and place the filter tube in the same collection tube.

19 Add 500 µL of Roche Washing Buffer to the filter tube.

20 Discard the flow-through and place the filter tube in the same collection tube.

> **Note**
>
> Ethanol removes remaining salts including leftover guanidium chloride.

21 Centrifuge the tube at 8,000 RCF for 1 minute.

22 Discard the flow-through and place the filter tube in the same collection tube.

23 Centrifuge the tube for 10 seconds at maximum speed to remove any residual ethanol, drying the filter column.

#### Elution of DNA

24 Transfer the filter tube to a 1.5 mL tube.

25 Add 200 µL of pre-warmed 55°C Roche Elution Buffer (pH 8.5 Tris-HCl).

> **Note**
>
> Re-hydrates the soluble DNA to allow it to flow through the filter. DNA is more stable and dissolves faster at this slightly basic pH than in water

26 Centrifuge the tube at 8,000 RCF for 1 minute to elute the DNA.

> **Note**
>
> The quantification of DNA concentrations (ng/µl) can be done using a Nanodrop spectrophotometer.

#### Genotyping PCR

27 The New England Biolabs (NEB) website can be consulted for the finer details of reaction volumes and components when amplifying using a standard NEB Taq DNA Polymerase and the 10X Standard Taq Reaction Buffer. The primers we used are listed in Supplementary Figure S2. Note that the annealing temperature is subject to variability according to the primer pair sequence. We recommend using 50-100 ng of genomic DNA as the template, as well as assembling all reaction components on ice prior to transferring to the preheated thermocycler. After the PCR, confirm the expected band sizes by carefully visualising the amplified products on a 1% agarose gel.

28 As a guide, the PCR program on the thermal cycler for a 1KB amplicon is as follows:

| A | B | C |
| --- | --- | --- |
| 1 cycle | 95°C | 2 minutes |
| 25 cycles | 95°C | 30 seconds |
| 60°C | 30 seconds | |
| 72°C | 60 seconds | |
| 1 cycle | 72°C | 5 minutes (to finish replication on all templates) |
| 1 cycle | 4-10°C | indefinite period (storing the sample prior to further analysis) |

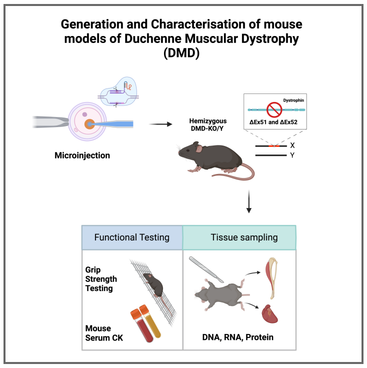


### Lateral Tail Vein Bleeding for Blood Collection in Mice

Jayshen Arudkumar[1], Yu Chinn Joshua Chey[1], Sandra Piltz[1], Paul Quinton Thomas[1], Fatwa Adikusuma[1]

[1]University of Adelaide

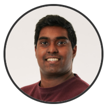

Jayshen Arudkumar
University of Adelaide, South Australian Health and Medical ...

#### DISCLAIMER

These protocols are for research purposes only.

#### ABSTRACT

**Paper abstract:** CRISPR-Cas9 gene-editing technology has revolutionised the creation of precise and permanent modifications to DNA, enabling the generation of diverse animal models for investigating potential treatments. Here, we provide a protocol for the use of CRISPR-Cas9 to create murine models of Duchenne Muscular Dystrophy (DMD) along with a step-by-step guide for their phenotypic and molecular characterisation. The experimental procedures include CRISPR microinjection of embryos, molecular testing at the DNA, RNA, and protein levels, forelimb grip strength testing, immunostaining and serum creatine kinase (CK) testing. We further provide suggestions for analysis and interpretation of the generated data, as well as the limitations of our approach. These protocols are designed for researchers who intend on generating and using mouse models to study DMD as well as those seeking a detailed framework of phenotyping to contribute to the broader landscape of genetic disorder investigations.
**Protocol summary:** Here we demonstrate the sampling of blood from the lateral tail vein of mice. Tail vein bleeding is necessary to isolate serum for downstream assessment of serum creatine kinase (CK) levels in our mouse model of Duchenne Muscular Dystrophy (DMD). We anticipate elevated serum CK levels in the DMD model compared to its wildtype counterparts, serving as a key indicator of the pathological changes associated with DMD.

#### IMAGE ATTRIBUTION

BioRender was used to generate figures for this manuscript.

**Protocol Info:** Jayshen Arudkumar, Yu Chinn Joshua Chey, Sandra Piltz, Paul Quinton Thomas, Fatwa Adikusuma . Lateral Tail Vein Bleeding for Blood Collection in Mice. **protocols.io** https://protocols.io/view/lateral-tail-vein-bleeding-for-blood-collection-in-c7yrzpv6

**Created:** Jan 23, 2024

**Last Modified:** Jan 30, 2024

**PROTOCOL integer ID:** 93937

**Keywords:** Serum, Mouse, Creatine Kinase, Phenotyping, DMD

MATERIALS

- Individually ventilated cages (IVC)
- PPE (Personal Protective Equipment) - gloves, scrubs, mop cap, and unit shoes
- Empty IVC cage with bedding
- 70% ethanol, F10 SC Veterinary Disinfectant
- Scalpel blade
- Non-treated Microvette® CB300 Blood collection System (Kent Scientific)
- EMLA cream (5%)
- Tissues
- Eppendorf 1.5 or 2 ml tubes
- Box (or Esky) and ice
- Marker pen
- Small gauze or swaddle

SAFETY WARNINGS

Wear proper PPE (gloves, safety goggles, enclosed shoes and lab coat) and prepare solvents in a chemical fume hood. Dispose used solvents or waste material in an appropriate biohazard waste containers.

ETHICS STATEMENT

Animal work described in this manuscript has been approved and conducted under the oversight of the Animal Ethics Committee of South Australian Health and Medical Research Institute (SAHMRI) and The University of Adelaide.

BEFORE START INSTRUCTIONS

Ensure there is suitable approval from your local Animal Ethics Committee for the maximum allowable blood volume to be taken per mice and that any operator is properly trained by their institution prior to commencing this protocol.

#### Preparation

1 Label microvette tubes with animal ID

2 Prepare ice and transport facilities

3 Arrange microvettes for easy access during bleeding

4 Prepare the empty IVC cage with bedding

#### Skin Numbing and Thermacage Preparation

5 Take the cage of animals and check to see if each mice present no health concerns.

6 Apply EMLA cream to the tail's incision area and leave mice to sit for 5-min.

7 Use Thermacage heat box and ensure it is warmed to 37°C.

8 Place the animal in the Thermacage and let them sit for no longer than 10-min.

9 Gently place the animal in the restrainer – holding on to its tail.

#### Tail Bleeding and Collection

10 Make a small incision over the lateral tail vein.

11 Encourage blood flow and collect into the Microvette tube.

12 Stop bleeding by applying gentle pressure using gauze and return animal to the empty IVC cage to sit for 5-min

13 If vein stops bleeding early, nick opposite vein if more blood required to be collected

14 Return the animal to its home cage and keep samples at room temperature

#### Serum Isolation

15 Ensure blood samples are left for 15-30-min from their last collection

16 Spin samples at 2000 RCF for 10-min at 4°C

17 Take colourless serum off the blood clot and place in microcentrifuge tube.

18 Freeze serum in -80°C.

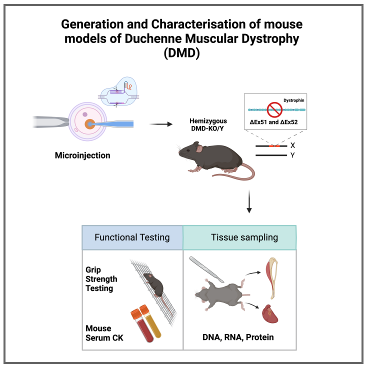


### Postmortem Mouse Tissue Processing (Necropsy)

Jayshen Arudkumar[1,2], Yu Chinn Joshua Chey[1,2], Sandra Piltz[1,2], Paul Quinton Thomas[1,2], Fatwa Adikusuma[1,2]

[1]University of Adelaide; [2]SAHMRI

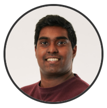

Jayshen Arudkumar
University of Adelaide, South Australian Health and Medical ...

#### DISCLAIMER

These protocols are for research purposes only.

#### ABSTRACT

**Paper abstract:** CRISPR-Cas9 gene-editing technology has revolutionised the creation of precise and permanent modifications to DNA, enabling the generation of diverse animal models for investigating potential treatments. Here, we provide a protocol for the use of CRISPR-Cas9 to create murine models of Duchenne Muscular Dystrophy (DMD) along with a step-by-step guide for their phenotypic and molecular characterisation. The experimental procedures include CRISPR microinjection of embryos, molecular testing at the DNA, RNA, and protein levels, forelimb grip strength testing, immunostaining and serum creatine kinase (CK) testing. We further provide suggestions for analysis and interpretation of the generated data, as well as the limitations of our approach. These protocols are designed for researchers who intend on generating and using mouse models to study DMD as well as those seeking a detailed framework of phenotyping to contribute to the broader landscape of genetic disorder investigations.
**Protocol summary:** A template for the steps preceding downstream molecular analyses of heart and skeletal muscle tissues. All procedures should be conducted in accordance with the ethical guidelines outlined by your animal ethics committee in your local institution.

#### IMAGE ATTRIBUTION

BioRender was used to generate figures for this manuscript.

**Protocol Info:** Jayshen Arudkumar, Yu Chinn Joshua Chey, Sandra Piltz, Paul Quinton Thomas, Fatwa Adikusuma . Postmortem Mouse Tissue Processing (Necropsy). **protocols.io** https://protocols.io/view/postmortem-mouse-tissue-processing-necropsy-c7ywzpxe

**Created:** Jan 23, 2024

**Last Modified:** Jan 30, 2024

**PROTOCOL integer ID:** 93942

**Keywords:** Dystrophin, DMD, Necropsy, Dissection, Mouse, Model

MATERIALS

- $CO_2$ chamber.
- Liquid nitrogen ($N_2$).
- Isopentane, isopentane container.
- 70% ethanol.
- Dissection tools (i.e., scalpels, scissors, tweezers, large forceps).
- Cork disc for cryostat use (20 mm x 3 mm), cork holder.
- Gum tragacanth (Sigma-Aldrich

SAFETY WARNINGS

Wear proper PPE (gloves, safety goggles, enclosed shoes and lab coat) and prepare solvents in a chemical fume hood. Dispose used solvents or waste material in an appropriate biohazard waste containers.

ETHICS STATEMENT

Animal work described in this manuscript has been approved and conducted under the oversight of the Animal Ethics Committee of South Australian Health and Medical Research Institute (SAHMRI) and The University of Adelaide.

#### Tissue Harvest and Freezing of Muscle Tissues

1 Euthanise the mouse using an AEC-approved method. Sacrifice mouse by CO2 exposure followed by cervical dislocation

2 From each mouse, we usually isolate the quadriceps, triceps, heart and tibialis anterior.

3 For downstream molecular analysis (DNA, RNA and Protein), cut a small portion of each isolated tissue while the remaining tissues will be subsequently frozen in liquid N2-cooled isopentane for cryo-sectioning.

**Note**

Ensure the muscle is in its normal physiological orientation without stretching.

4 Prepare the gum tragacanth mixture by mixing 10% (w/v) into water in a small beaker. Continue stirring and leave to set until a slurry forms.

5 Pour isopentane in a metal container and place within a larger container containing liquid nitrogen. Chill isopentane in liquid nitrogen until white solid precipitate is visible at the bottom of the container.

Note

The optimal temperature range for isopentane is around -150°C. Once within this temperature range, solid white pebbles of frozen isopentane will form at the bottom. Avoid freezing artifacts by ensuring freezing occurs after this stage. In the event that the isopentane freezes solid, allow it to thaw and then re-chill to freezing temperatures before using it again.

6 Place a conservative amount of slurry on the cork pads suitable for tissue mounting. Mount the tissue sections on to the cork to ensure at least 2/3rds of the section remain upright and oriented for a transverse section.

7 Using pre-chilled forceps, lower the specimen and cork into the isopentane and suspend for the appropriate time (20s for smaller tissues, 30s for heart and quad), depending on muscle size.

8 Transfer specimen immediately to dry ice and label to store in a -80°C freezer until sectioning.

#### Prepare Tissue Block for Cutting

9 Transport tissues on dry ice to the cryostat to prevent thawing.

10 Set the cryostat chamber to -20°C to -24°C and let frozen samples equilibrate for 30 min.

11 Apply a uniform thin layer of OCT to the specimen disc. With the tissue freeze mounted on the cork, place the sample at the correct orientation.

#### Sectioning of Tissue

12 Mount the specimen onto the specimen head in the cryostat, ensuring good contact.

13 Set the chamber temperature to -21°C to -24°C and the chamber head temperature to -15°C.

14 Adjust thickness settings (7-10 µm) and trim the block as needed.

15 Cut sections perpendicular to the direction of myofiber orientation.

16 Collect sections on pre-warmed positively charged microscope slides by pressing them onto the sections.

17 Inventory and archive slides at -20°C for long-term storage or proceed to staining.

### Scientific Protocol for Immunofluorescence Analysis: Dystrophin

Jayshen Arudkumar[1,2], Yu Chinn Joshua Chey[1,2], Sandra Piltz[1,2], Paul Quinton Thomas[1,2], Fatwa Adikusuma[1,2]

[1]University of Adelaide; [2]SAHMRI

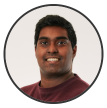

Jayshen Arudkumar
University of Adelaide, South Australian Health and Medical ...

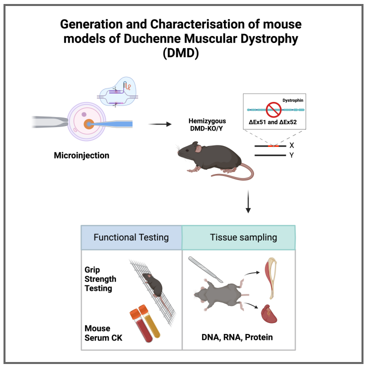


**Protocol Info:** Jayshen Arudkumar, Yu Chinn Joshua Chey, Sandra Piltz, Paul Quinton Thomas, Fatwa Adikusuma . Scientific Protocol for Immunofluorescence Analysis: Dystrophin. **protocols.io** https://protocols.io/view/scientific-protocol-for-immunofluorescence-analysi-c7yyzpxw

**Created:** Jan 23, 2024

**Last Modified:** Jan 30, 2024

**PROTOCOL integer ID:** 93944

**Keywords:** Immunofluorescence, DMD, Dystrophin, Phenotyping, Mouse, Knockout

#### DISCLAIMER

These protocols are for research purposes only.

#### ABSTRACT

**Paper abstract:** CRISPR-Cas9 gene-editing technology has revolutionised the creation of precise and permanent modifications to DNA, enabling the generation of diverse animal models for investigating potential treatments. Here, we provide a protocol for the use of CRISPR-Cas9 to create murine models of Duchenne Muscular Dystrophy (DMD) along with a step-by-step guide for their phenotypic and molecular characterisation. The experimental procedures include CRISPR microinjection of embryos, molecular testing at the DNA, RNA, and protein levels, forelimb grip strength testing, immunostaining and serum creatine kinase (CK) testing. We further provide suggestions for analysis and interpretation of the generated data, as well as the limitations of our approach. These protocols are designed for researchers who intend on generating and using mouse models to study DMD as well as those seeking a detailed framework of phenotyping to contribute to the broader landscape of genetic disorder investigations.
**Protocol summary:** Here we detect dystrophin on the muscle membrane of heart and skeletal muscle tissues. These include sections from the heart, tibialis anterior (TA), triceps and quadriceps. Antibodies used for immunofluorescence can be found in the linked publication, in Supplementary Table S3.

#### IMAGE ATTRIBUTION

BioRender was used to generate figures for this manuscript.

MATERIALS

- SuperfrostTM slides (ThermoFisher)
- Cryostat machine
- PPE (Personal Protective Equipment)
- Triton X-100 (X100, Sigma-Aldrich)
- Phosphate-buffered saline 1x (PBS)
- Kimwipe Tissues
- Eppendorf 1.5 or 2 ml tubes
- Box (or Esky) and ice
- ReadyProbes**TM** Hydrophobic Barrier Pap Pen (Abcam)
- Coverslip
- ProLong**TM** Gold Antifade Mountant with DNA Stain DAPI (ThermoFisher)
- ReadyProbes**TM** Mouse-on-Mouse IgG Blocking Solution (ThermoFisher)
- Fetal Bovine Serum (Gibco)

SAFETY WARNINGS

Wear proper PPE (gloves, safety goggles, enclosed shoes and lab coat) and prepare solvents in a chemical fume hood. Dispose used solvents or waste material in an appropriate biohazard waste containers.

ETHICS STATEMENT

Animal work described in this manuscript has been approved and conducted under the oversight of the Animal Ethics Committee of South Australian Health and Medical Research Institute (SAHMRI) and The University of Adelaide.

#### Tissue Harvesting and Preparation

1 For tissue isolation and freezing of muscle tissues using liquid nitrogen samples snap frozen on gum tragacanth-mounted corkpads, see the previous protocol titled ‘Postmortem Tissue Processing’. The frozen tissues should be stored in the -80°C until ready for sectioning.

#### Preparation of Slides

2 Allow the slides to air dry for 5 minutes to eliminate any residual moisture.

3 Ensure the absence of moisture before initiating immunostaining procedures.

#### Primary Antibody Stain + Blocking

4 Cover the tissue sections with 4% paraformaldehyde (PFA) or ice-cold methanol and incubate for 15 minutes.

5 Aspirate the PFA mixture and wash the tissue sections gently with phosphate-buffered saline (PBS).

6 Create a hydrophobic barrier around the tissue sections using a PAP pen.

7 Incubate the slides in a permeabilization buffer (0.3% PBS-Triton X-100) for 10 minutes inside a humidified chamber.

8 Cover the slides with a blocking solution (0.3% PBS-Tween + 10% fetal bovine serum, FBS) and incubate for at least 1 hour at room temperature.

9 Cover the slides with 1x Thermo ReadyProbe MOM solution and incubate for 1 hour at room temperature (use 1 drop in 1.25 mL PBS).

10 Wash the slides three times with PBS for 5 minutes each.

11 Dilute primary antibodies in blocking solution (e.g., Mandys8 at 1:100). Incubate the slides overnight at 4°C.

#### Secondary Antibody Stain + Imaging

12 Wash the slides three times with PBS for 5 minutes each.

13 Cover the slides with secondary antibodies diluted in blocking buffer (e.g., 1:300 Anti-Mouse 594) for 1 hour at room temperature in the dark. Keep slides in the dark for the remainder of the protocol.

14 Wash the slides three times with PBS for 5 minutes each.

#### Mounting and Storage

15 Mount a coverslip on the slides using a mounting medium such as ProLong™ Gold Antifade Mountant with DNA Stain DAPI

16 Allow the slides to air dry in the dark for 1–2 hours.

17 Store the slides at either 4°C or -20°C and visualize within 2 weeks.

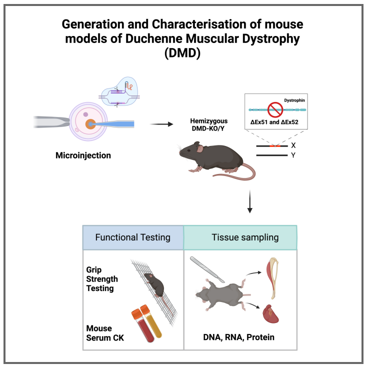


**Protocol Info:** Jayshen Arudkumar, Yu Chinn Joshua Chey, Sandra Piltz, Paul Quinton Thomas, Fatwa Adikusuma . Scientific Protocol for H&E staining in Heart and Skeletal Muscle. **protocols.io** https://protocols.io/view/scientific-protocol-for-h-amp-e-staining-in-heart-c7yzzpx6

**Created:** Jan 23, 2024

**Last Modified:** Jan 30, 2024

**PROTOCOL integer ID:** 93945

**Keywords:** H&E, Histology, CRISPR, DMD, Muscle, Heart

### Scientific Protocol for H&E staining in Heart and Skeletal Muscle

Jayshen Arudkumar[1,2], Yu Chinn Joshua Chey[1,2], Sandra Piltz[1,2], Paul Quinton Thomas[1,2], Fatwa Adikusuma[1,2]

[1]University of Adelaide; [2]SAHMRI

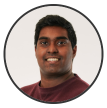

Jayshen Arudkumar
University of Adelaide, South Australian Health and Medical ...

#### DISCLAIMER

These protocols are for research purposes only.

#### ABSTRACT

**Paper abstract:** CRISPR-Cas9 gene-editing technology has revolutionised the creation of precise and permanent modifications to DNA, enabling the generation of diverse animal models for investigating potential treatments. Here, we provide a protocol for the use of CRISPR-Cas9 to create murine models of Duchenne Muscular Dystrophy (DMD) along with a step-by-step guide for their phenotypic and molecular characterisation. The experimental procedures include CRISPR microinjection of embryos, molecular testing at the DNA, RNA, and protein levels, forelimb grip strength testing, immunostaining and serum creatine kinase (CK) testing. We further provide suggestions for analysis and interpretation of the generated data, as well as the limitations of our approach. These protocols are designed for researchers who intend on generating and using mouse models to study DMD as well as those seeking a detailed framework of phenotyping to contribute to the broader landscape of genetic disorder investigations.
**Protocol summary:** The following protocol describes H&E staining of snap frozen cork-mounted mouse tissue samples. These include sections from the heart, tibialis anterior (TA), triceps and quadriceps.

#### IMAGE ATTRIBUTION

BioRender was used to generate figures for this manuscript.

MATERIALS

- Lillie Mayer's Haematoxylin (0.5%)
- Tap water
- 0.1% Eosin
- Coverslip
- DPX mountant (Sigma-Aldrich)
- Slide staining tray, rack (ProSci Tech)

SAFETY WARNINGS

Wear proper PPE (gloves, safety goggles, enclosed shoes and lab coat) and prepare solvents in a chemical fume hood. Dispose used solvents or waste material in an appropriate biohazard waste containers.

ETHICS STATEMENT

Animal work described in this manuscript has been approved and conducted under the oversight of the Animal Ethics Committee of South Australian Health and Medical Research Institute (SAHMRI) and The University of Adelaide.

#### Preparation of Slides

1 Allow the slides to air dry for 5 minutes to eliminate any residual moisture.

#### Haematoxylin & Eosin Staining

2 Immerse the slide in the following reagents order:

2.1 Haematoxylin for 5 min. Drain the excess of the hematoxylin by tapping the slide tray on paper towels

2.2 Place slide staining tray into a container and run through tap water for 2-minutes

2.3 Eosin stain solution for 3 min. Drain the excess eosin by tapping the slide tray on paper towels

#### Dehydration Steps

2.4 Dip slide tray twice into 70% ethanol

2.5 Dip slide tray four times into 95% ethanol

2.6 100% ethanol for 1 min and repeat in clean 100% ethanol for 1 min.

2.7 Xylene for 1 min and repeat in clean xylenes for 1 min. Drain the excess xylenes by tapping the slide on paper towels.

#### Mounting and Imaging

3 Add one drop of DPX mounting medium to the tissue section and cover with an appropriate-sized coverslip. Drain any excess mounting media. Allow the slide to air dry in a fume hood.

4 Visualise the stained tissue under a light microscope using bright-field illumination.

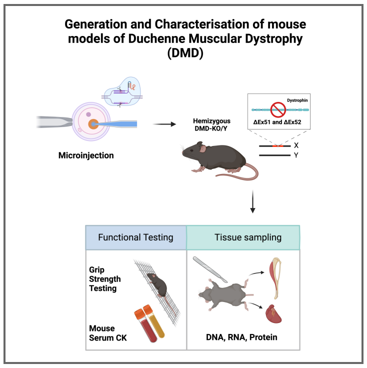


### RNA Extraction and RT-PCR on Mouse tissues

Jayshen Arudkumar[1,2], Yu Chinn Joshua Chey[1,2], Sandra Piltz[1,2], Paul Quinton Thomas[1,2], Fatwa Adikusuma[1,2]

[1]University of Adelaide; [2]SAHMRI

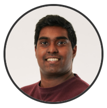

Jayshen Arudkumar
University of Adelaide, South Australian Health and Medical ...

#### DISCLAIMER

These protocols are for research purposes only.

**Protocol Info:** Jayshen Arudkumar, Yu Chinn Joshua Chey, Sandra Piltz, Paul Quinton Thomas, Fatwa Adikusuma . RNA Extraction and RT-PCR on Mouse tissues. **protocols.io** https://protocols.io/view/rna-extraction-and-rt-pcr-on-mouse-tissues-c732zqqe

**Created:** Jan 24, 2024

**Last Modified:** Jan 30, 2024

**PROTOCOL integer ID:** 94042

**Keywords:** RNA, RT-PCR, Dystrophin, transcript, Phenotyping, Knockout, DMD

#### ABSTRACT

**Paper abstract:** CRISPR-Cas9 gene-editing technology has revolutionised the creation of precise and permanent modifications to DNA, enabling the generation of diverse animal models for investigating potential treatments. Here, we provide a protocol for the use of CRISPR-Cas9 to create murine models of Duchenne Muscular Dystrophy (DMD) along with a step-by-step guide for their phenotypic and molecular characterisation. The experimental procedures include CRISPR microinjection of embryos, molecular testing at the DNA, RNA, and protein levels, forelimb grip strength testing, immunostaining and serum creatine kinase (CK) testing. We further provide suggestions for analysis and interpretation of the generated data, as well as the limitations of our approach. These protocols are designed for researchers who intend on generating and using mouse models to study DMD as well as those seeking a detailed framework of phenotyping to contribute to the broader landscape of genetic disorder investigations.
**Protocol summary:** To obtain high-quality RNA for subsequent molecular analyses, specifically to assess mRNA transcript expression in mouse tissues. Following this, cDNA synthesis is employed to convert the isolated RNA into complementary DNA (cDNA). This cDNA serves as a stable template for downstream analyses, where we look at using reverse transcription PCR (RT-PCR) or quantitative real-time PCR (qPCR).

#### IMAGE ATTRIBUTION

BioRender was used to generate figures for this manuscript.

MATERIALS

- TRIzolTM Reagent (Invitrogen).
- Chloroform.
- MagNA Lyser Green Beads (Roche)
- Ethanol.
- Isopropanol.
- DNase/RNase-free distilled water.
- RNEasy Mini Extraction kit (Qiagen)
- High-Capacity cDNA Reverse Transcription Kit (Applied Biosystems)
- QuantStudio Real-Time PCR software v1.3 (Applied Biosystems)

SAFETY WARNINGS

Wear proper PPE (gloves, safety goggles, enclosed shoes and lab coat) and prepare solvents in a chemical fume hood. Dispose used solvents or waste material in an appropriate biohazard waste containers.

ETHICS STATEMENT

Animal work described in this manuscript has been approved and conducted under the oversight of the Animal Ethics Committee of South Australian Health and Medical Research Institute (SAHMRI) and The University of Adelaide.

#### Preparation Phase

1 Keep the tissue samples on ice

2 Set the centrifuge to cool at 4°C

3 Mince the heart and skeletal muscle tissues in a few drops of water and place the minced portions into a magNA lyser tube for bead homogenisation step

4 Homogenise samples in the Precellys bead-based homogeniser at 6500 RPM for 2 cycles at 20s each

#### TRIzol Treatment

5 Add 500 µL of Invitrogen TRIzol reagent to the supernatant

6 Incubate for 5 minutes at room temperature (RT) to dissociate the nucleoprotein complex

#### Chloroform Extraction

7 Add 100 µL chloroform

**Note**

Use 0.2 ml chloroform/ 1 ml TRIZOL reagent

8 Shake the tubes vigorously by hand for 15 seconds to mix

9 Incubate for 2 min at RT.

10 Centrifuge at 12,000 RCF in a 4°C cooled centrifuge for 15 minutes.

#### RNA Precipitation and Collection

11 Label new microcentrifuge tubes and keep on ice

12 Transfer approximately 175 µL of the colourless liquid layer to a new eppendorf tube.

13 Add an equal volume of 70% ethanol. Mix by pipetting. Invert the tube and mix to disperse any visible precipitate that can form after adding ethanol.

14 Transfer a maximum of 700 µL to the QIAGEN spin column with a collection tube.

15 Centrifuge for 30 seconds at 8,000 RCF (room temperature centrifuge).

16 Place the flow-through back into the spin column and centrifuge again for 30 seconds at 6,500 RCF. Discard the flow-through and place the column back into the collection tube.

17 Add 700 µL Buffer RW1 to the RNeasy spin column. Centrifuge for 15 seconds at 8,000 RCF.

18 Add 500 µL Buffer RPE to the RNeasy spin column. Centrifuge for 15 seconds at 8,000 RCF.

19 Discard the flow-through and place the spin column in a new collection tube.

20 Add 500 µL Buffer RPE to the RNeasy spin column. Centrifuge for 2 minutes at 8,000 RCF to wash the spin column membrane.

21 Discard the flow-through and place the spin column in a new collection tube.

22 Place the RNeasy spin column in a new collection tube. Centrifuge for 1 minute at max speed.

23 Place the RNeasy spin column in a new QIAGEN RNase-free 1.5 ml tube.

24 Add 30-50 µL RNase-free water directly to the spin column membrane.

25 Centrifuge for 1 minute at 8,000 RCF to elute the RNA.

Note

For <100 µg starting tissue, use one 30-100 µL elution volume. For >100 µg, use 2–3 sequential 100 µL elutions.

26 Measure concentration of the RNA solution using NanoDrop Spectrophotometer

Note

The A260/A280 ratio should be approximately 2.0, but values between 1.8 and 2.1 are considered to be of acceptable purity.

#### Reverse Transcription Reaction for cDNA Generation

27 Follow the protocol provided by the High-Capacity cDNA Reverse Transcription Kit (Applied Biosystems) to convert 1-2 µg of RNA into cDNA

28 Dilute cDNA in 50 µL ultrapure water prior to running downstream analysis

#### Reverse-transcription polymerase chain reaction (RT-PCR)

29 The New England Biolabs (NEB) website can be consulted for the finer details of reaction volumes and components when amplifying using a standard NEB Taq DNA Polymerase and the 10X Standard Taq Reaction Buffer. The primers we used are listed in Supplementary Figure S2. We recommend using cDNA synthesised from 1-2 µg of RNA, as well as assembling all rection components on ice prior to transferring to the preheated thermocycler. After the RT-PCR, confirm the expected band sizes by carefully visualizing the amplified products on a 1% agarose gel.

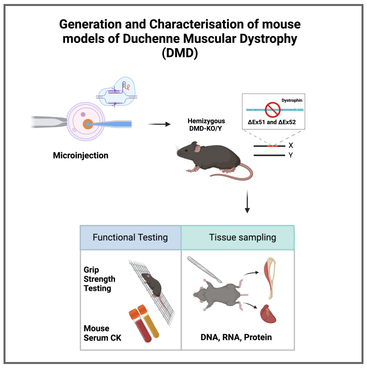


### Scientific Protocol for Western Blotting

Jayshen Arudkumar[1,2], Yu Chinn Joshua Chey[1,2], Sandra Piltz[1,2], Paul Quinton Thomas[1,2], Fatwa Adikusuma[1,2]

[1]University of Adelaide; [2]SAHMRI

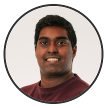

Jayshen Arudkumar
University of Adelaide, South Australian Health and Medical ...

#### DISCLAIMER

These protocols are for research purposes only.

**Protocol Info:** Jayshen Arudkumar, Yu Chinn Joshua Chey, Sandra Piltz, Paul Quinton Thomas, Fatwa Adikusuma . Scientific Protocol for Western Blotting. **protocols.io** https://protocols.io/view/scientific-protocol-for-western-blotting-c736zqre

**Created:** Jan 24, 2024

**Last Modified:** Jan 30, 2024

**PROTOCOL integer ID:** 94046

**Keywords:** Protein, CRISPR, DMD, Mouse, PVDF, Blot, Western, Knockout

#### ABSTRACT

**Paper abstract:** CRISPR-Cas9 gene-editing technology has revolutionised the creation of precise and permanent modifications to DNA, enabling the generation of diverse animal models for investigating potential treatments. Here, we provide a protocol for the use of CRISPR-Cas9 to create murine models of Duchenne Muscular Dystrophy (DMD) along with a step-by-step guide for their phenotypic and molecular characterisation. The experimental procedures include CRISPR microinjection of embryos, molecular testing at the DNA, RNA, and protein levels, forelimb grip strength testing, immunostaining and serum creatine kinase (CK) testing. We further provide suggestions for analysis and interpretation of the generated data, as well as the limitations of our approach. These protocols are designed for researchers who intend on generating and using mouse models to study DMD as well as those seeking a detailed framework of phenotyping to contribute to the broader landscape of genetic disorder investigations.
**Paper summary:** The following protocol describes the separation of proteins through SDS-PAGE, the transfer onto a PVDF membrane and subsequent staining with dystrophin-specific antibodies.

#### IMAGE ATTRIBUTION

BioRender was used to generate figures for this manuscript.

MATERIALS

- Complete™, Mini, EDTA-free Protease Inhibitor Cocktail tablet (Roche)
- Pierce™ BCA Protein Assay Kit (ThermoFisher)
- MagNA Lyser Green Beads (Roche)
- Pierce™ Protease Inhibitor Mini Tablets, EDTA-free (ThermoFisher)
- 96-well plates
- SureLock™ Tandem Midi Gel Tank (Invitrogen)
- Trans-Blot Turbo™ Transfer System (Bio-Rad)
- Sample loading buffer: 75 mM Tris–HCl at pH 6.8, 5 mM EDTA, 15% SDS, 20% glycerol, 0.004% w/v bromophenol blue, and 5% β-mercaptoethanol
- NuPAGE™ NovexTM 3–8% Tris–Acetate Midi 20-well gel (Invitrogen)
- NuPAGE™ Tris–Acetate SDS running buffer (Invitrogen)
- HiMark™ pre-stained protein standard (ThermoFisher)
- Trans-Blot Turbo Midi 0.2 µm PVDF Transfer Packs, 0.20 µm pore size (Bio-Rad).
- Wash Buffer: PBST (phosphate-buffered saline, 0.1% Tween 20)
- Blocking: PBST and 5% skimmed milk
- Primary antibodies: MANDYS8 Monoclonal Ant-Dystrophin antibody (Sigma-Aldrich) and V9131 Anti-Vinculin antibody (Sigma-Aldrich)
- Secondary antibodies: Goat anti-mouse immunoglobulin G (IgG) (H+L) secondary antibody, horseradish peroxidase (HRP) (ThermoFisher).
- Supersignal West Femto kit (ThermoFisher)
- Blot Shaker
- Skim milk powder (Diploma)
- Blotting Tweezer (Invitrogen)
- Blot roller (Bio-Rad)
- Heat block
- Centrifuge
- Incubation Tray 10 x 14 cm (ThermoFisher)
- Bio-Rad ChemiDoc MP Imaging System
- Restore™ PLUS Western Blot Stripping Buffer (ThermoFisher)

SAFETY WARNINGS

Wear proper PPE (gloves, safety goggles, enclosed shoes and lab coat) and prepare solvents in a chemical fume hood. Dispose used solvents or waste material in an appropriate biohazard waste containers.

ETHICS STATEMENT

Animal work described in this manuscript has been approved and conducted under the oversight of the Animal Ethics Committee of South Australian Health and Medical Research Institute (SAHMRI) and The University of Adelaide.

#### Protein Extraction and BCA

1 Transfer isolated skeletal muscle and heart tissues to magna lyser bead tubes

2 Add 480 µL Disruption Buffer (15% SDS, 75 mM Tris HCl pH 6.8) + 20 µL 25x PI cocktail (optional).

3 Homogenise samples in the Precellys bead-based homogeniser at 6500 RPM for 2 cycles 20 s each

4 Spin down the supernatant in a 4°C cooled centrifuge at 13,500 RCF for 10 minutes

5 Collect the supernatant and place into a new 1.5 ml tube

6 Perform BCA Assay and quantitation steps according to manufacturer’s protocol, using the disruption buffer as a blank.

Note

25 µg of total protein will be used from each tissue per run

7 Store protein sample as multiple low volume aliquots at -80°C until needed for SDS-PAGE

#### SDS-PAGE

8 Load equal parts of sample loading buffer with sample to make up 25 µg protein

Note

We use a 20-combed 3-8% Polyacrylamide gel that is pre-cast. The maximum load volume in each well is 25 µl.

9 Denature samples at 98°C for 5 minutes

Note

While samples are left on heating block, prepare the gel tank with midi-well immersed in running buffer. Ensure the buffer is above the electrode line.

10 Briefly centrifuge samples to collect the mixture

11 Load samples into the gel and run the gel at 100 V for 15 minutes, followed by 120 V for 1 hour and 45 minutes.

Note

In alignment with the recommendation from ThermoFisher, we have also found that running at 150 V for 1h worked as effectively.

#### Semi-dry transfer (TransBlot Turbo System) and Block

12 Transfer the protein from the gel to the membrane and place into Trans-Blot Turbo Transfer System cassette.

Note

Ensure that the PVDF membrane is kept moist on the top and bottom filter paper stack. Once gel is neatly stacked on the membrane, firmly roll out the air bubbles with a roller.

13 Ensure that the whole transfer stack is firm when closing the cassette lid to ensure no additional air bubbles form

14 Place transfer stack inside the Trans-Blot Turbo Transfer System machine and run using the 'HIGH MW' protocol in the standard Bio-Rad setting.

15 Place the membrane in a blotting container and add enough blocking solution (10% Skim milk in PBST) to cover the entire membrane surface

Note

Recommended to pour solutions gently into the corner of the blotting container and not directly onto the membrane so that the proteins on the membranes are not disturbed.

16 Incubate the membrane on a blot shaker at 50 – 60 RPM for 1h

17 Pour out the blocking solution

#### Primary and Secondary Antibody Incubations

18 Incubate the membrane with primary antibody at a dilution of 1:1000 (MANDYS8) in blocking buffer (2% Skim milk in PBST), for an overnight incubation at 4°C

19 Wash the membrane in three washes of PBST on a blot shaker for 5 min each

20 Incubate the membrane with HRP-conjugated secondary antibody (Goat anti-mouse IgG HRP) in blocking buffer (2% Skim milk in PBST) at a dilution of 1:8000, for 1 hour at room temperature.

21 Wash the membrane in three washes of PBST on a blot shaker for 5 min each

#### Detection and Imaging

22 Detect the signal following the Supersignal West Femto Maximum Sensitivity Substrate kit recommendations

**Note**

For a Midi (8.5 x 13.5 cm) membrane, we use 2 ml of West Femto Working Solution mix. In a yellow-capped tube, mix equal parts of West Femto Stable Peroxide agent (1 mL) and West Femto Luminol/Enhancer reagent (1 mL). This Working Solution forms the substrate for chemiluminescence. Pipette enough of the substrate onto a parafilm paper. Incubate the membrane under dark conditions for 5-min before removing excess reagent.

23 Arrange the membrane on the imaging surface, taking care not to create bubbles on the surface.

24 Image the blot using your machine of choice

**Note**

We use the Bio-Rad Chemidoc MP Imaging system using the Chemi Hi Sensitivity setting. The exposure time can be manually adjusted to obtained the best image prior to saturation. This system enables you to take a multichannel image with the colorimetric and either the Chemi Hi Sensitivity or Hi-Resolution setting to get an image of the chemiluminescent dystrophin bands overlaid with the protein size marker – evident from the colorimetric setting. Verify the molecular weight of the band of interest (should be close to 460kDa mark) based on the multichannel image.

#### Stripping and Re-probe with Vinculin

25 Strip the membrane using the Restore PLUS Western Stripping Buffer, following manufacturer's instructions. This includes a membrane washing and subsequent blocking step prior to re-probing.

26 Re-probe by incubating the membrane with vinculin-targeting primary antibody at a dilution of 1:1000 (V9131) in blocking buffer (2% Skim milk in PBST) at room temperature for 1h.

27 Follow steps 17-22 until imaging of the blot

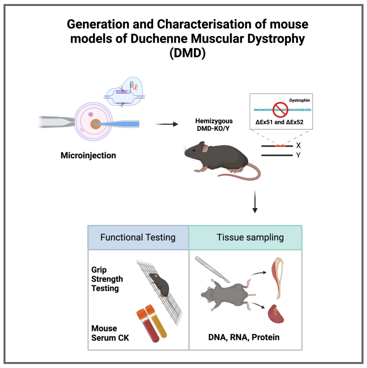


### Forelimb Grip Strength testing

Jayshen Arudkumar[1,2], Yu Chinn Joshua Chey[1,2], Sandra Piltz[1,2], Paul Quinton Thomas[1,2], Fatwa Adikusuma[1,2]

[1]University of Adelaide; [2]SAHMRI

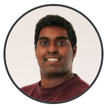

Jayshen Arudkumar
University of Adelaide, South Australian Health and Medical ...

#### DISCLAIMER

These protocols are for research purposes only.

**Protocol Info:** Jayshen Arudkumar, Yu Chinn Joshua Chey, Sandra Piltz, Paul Quinton Thomas, Fatwa Adikusuma . Forelimb Grip Strength testing. **protocols.io**
https://protocols.io/view/forelimb-grip-strength-testing-c737zqrn

**Created:** Jan 24, 2024

**Last Modified:** Jan 30, 2024

**PROTOCOL integer ID:** 94047

**Keywords:** CRISPR, Grip strength, DMD, Phenotyping, Mice

#### ABSTRACT

**Paper abstract:** CRISPR-Cas9 gene-editing technology has revolutionised the creation of precise and permanent modifications to DNA, enabling the generation of diverse animal models for investigating potential treatments. Here, we provide a protocol for the use of CRISPR-Cas9 to create murine models of Duchenne Muscular Dystrophy (DMD) along with a step-by-step guide for their phenotypic and molecular characterisation. The experimental procedures include CRISPR microinjection of embryos, molecular testing at the DNA, RNA, and protein levels, forelimb grip strength testing, immunostaining and serum creatine kinase (CK) testing. We further provide suggestions for analysis and interpretation of the generated data, as well as the limitations of our approach. These protocols are designed for researchers who intend on generating and using mouse models to study DMD as well as those seeking a detailed framework of phenotyping to contribute to the broader landscape of genetic disorder investigations.
**Protocol summary:** We use the forelimb grip strength test to determine the maximum force applied by the mouse onto a grid, as an evaluation of the muscle integrity in the DMD models. The test is based on the tendency of a mouse to instinctively grasp a grid when suspended by the tail, where there is a measure of the maximal peak force generated from the combined front paws (1, 2). The protocol listed here was adapted from the TREAT-NMD standard operating procedure SOP DMD_M.2.2.001 for grip strength testing.

#### IMAGE ATTRIBUTION

BioRender was used to generate figures for this manuscript.

#### MATERIALS

- Weighing scale
- Andilog Force Gauge Centor Easy 250N
- Metal grid (SDR Scientific)
- F10 SC Veterinary Disinfectant

#### SAFETY WARNINGS

Wear proper PPE (gloves, safety goggles, enclosed shoes and lab coat) and prepare solvents in a chemical fume hood. Dispose used solvents or waste material in an appropriate biohazard waste containers.

#### ETHICS STATEMENT

Animal work described in this manuscript has been approved and conducted under the oversight of the Animal Ethics Committee of South Australian Health and Medical Research Institute (SAHMRI) and The University of Adelaide.

### Apparatus Setup

1 We attached the grasping grid to the force gauge, using a hooked connector that extends from the base end of the gauge. The grasping grid accessory is a stainless-steel attachment that allows the mouse to grip with both front paws. With regards to the measurement settings, make sure the setting is on MAX and that the units of force is adjusted to grams-of-force.

### Testing

2 Prior to the test, weigh each mouse and return to their cage

3 Gently lower the mouse by the base of its tail and ensure that the mouse grasps the grid tightly with both front paws

4 Pull the mouse away from the grid so that its grasp is broken; the highest force applied to the grid will be shown on the force gauge's display. Manually record this value.

**Note**

Only take pulls into account in which the mouse shows resistance to the experimenter. Reject measures in which only one forepaw, or the hindlimbs were used and in which the mouse turned during the pull.

5 Reset the meter at the start of each recording

6 Let the mouse pull the grid three times in a row and then return it in the cage for a resting period of at least one minute.

**Note**

Between series of pulls a resting period is necessary for the mouse to recover and avoid habit formation.

7 Then let the mouse perform four series of pulls, each followed by a short 1-min resting period. In this way the mouse has pulled a total of 12x (3 pulls x 4 times = 12 total pulls).

8 Determine the maximum grip strength and normalize for body weight by taking the average of the three highest values out of the 12 values collected.
